# Tensor analysis of animal behavior by matricization and feature selection

**DOI:** 10.1101/2025.01.28.635088

**Authors:** Beichen Wang, Jiazhang Cai, Luyang Fang, Ping Ma, Yuk Fai Leung

## Abstract

Contemporary neurobehavior research often collects multi-dimensional tensor (MDT) data, consisting of time-series measurements for multiple features from multiple animals subjected to various perturbations. Proper analysis of the MDT data can facilitate the dissection of the underlying neural circuitry driving the behavior. However, many common approaches for MDT analysis, such as tensor decomposition, often yield results that are difficult to interpret and not directly compatible with standard multivariate analysis (MVA), which is designed for simpler, lower-dimensional data structures. To address this issue, dimensionality reduction techniques, including matricization methods such as Index Construction and Feature Concatenation, are applied to transform all or a subset of the features in the MDT into a lower-dimensional tensor, commonly a 2-dimensional tensor (2DT), that is compatible with MVA. However, the matricization methods may exclude information from the MDT features or create too many 2DT features that introduce spurious noise to the downstream analyses. Their impacts on the downstream MVA performance remain elusive. In this study, we systematically evaluated different approaches for matricization and feature selection and their impacts on MVA performance using an MDT dataset of zebrafish visual- motor response collected from wild-types (WTs) and visually-impaired mutants. We matricized the MDT dataset using various Index Construction and Feature Concatenation methods, then identified informative 2DT features using the filter and embedded methods. To evaluate these feature-selection approaches, we conducted a classification task distinguishing WT and visually-impaired zebrafish by multiple classifiers. We then assessed classification performance with cross-validation and holdout validation. We found that most classifiers performed the best when using all 2DT features matricized by Feature Concatenation and selected by the embedded method. The results also revealed unique behavioral differences between the WTs and visually-impaired mutants that were not identified by standard MVA or MDT analysis. Our results demonstrate the utility of analyzing MDT behavioral data by matricization and feature selection.

## Introduction

Animals display dynamic behaviors in response to environmental changes and stimulations, reflecting the underlying neural processing of their surroundings and experiences. Studying these animal behaviors facilitates the dissection of the neural processing at the cellular, circuitry, individual, and population levels [1–3]. The study approach can be observational [4] or experimental [5], wherein behavioral dynamics are recorded through one or more types of quantitative measurement [6,7]. Even when an experiment collects just one behavior measurement in a time series, e.g., the displacement of animals, the resulting dataset contains multiple samples and time points. Such data are regarded as multivariate data (Fig 1A) and typically analyzed by multivariate analysis (MVA), which includes multivariate hypothesis testing (e.g., Hotelling’s T^2^ Test [8], MANOVA [8], and PERMANOVA [9]) and machine learning (e.g., regression [10] and clustering [11,12]). As the variety of behavioral measurements increases with the advent of technology, a new issue emerges at the higher-level data structure, as illustrated in Fig 1B. In this data structure, even though each data point can be generally described by three factors or dimensions (color arrows in Fig 1B), each factor or dimension can contain multiple features (e.g., multiple distinct behavioral measurements, multiple samples with different genotypes and treatment conditions, or multiple time points).

**Fig 1.**
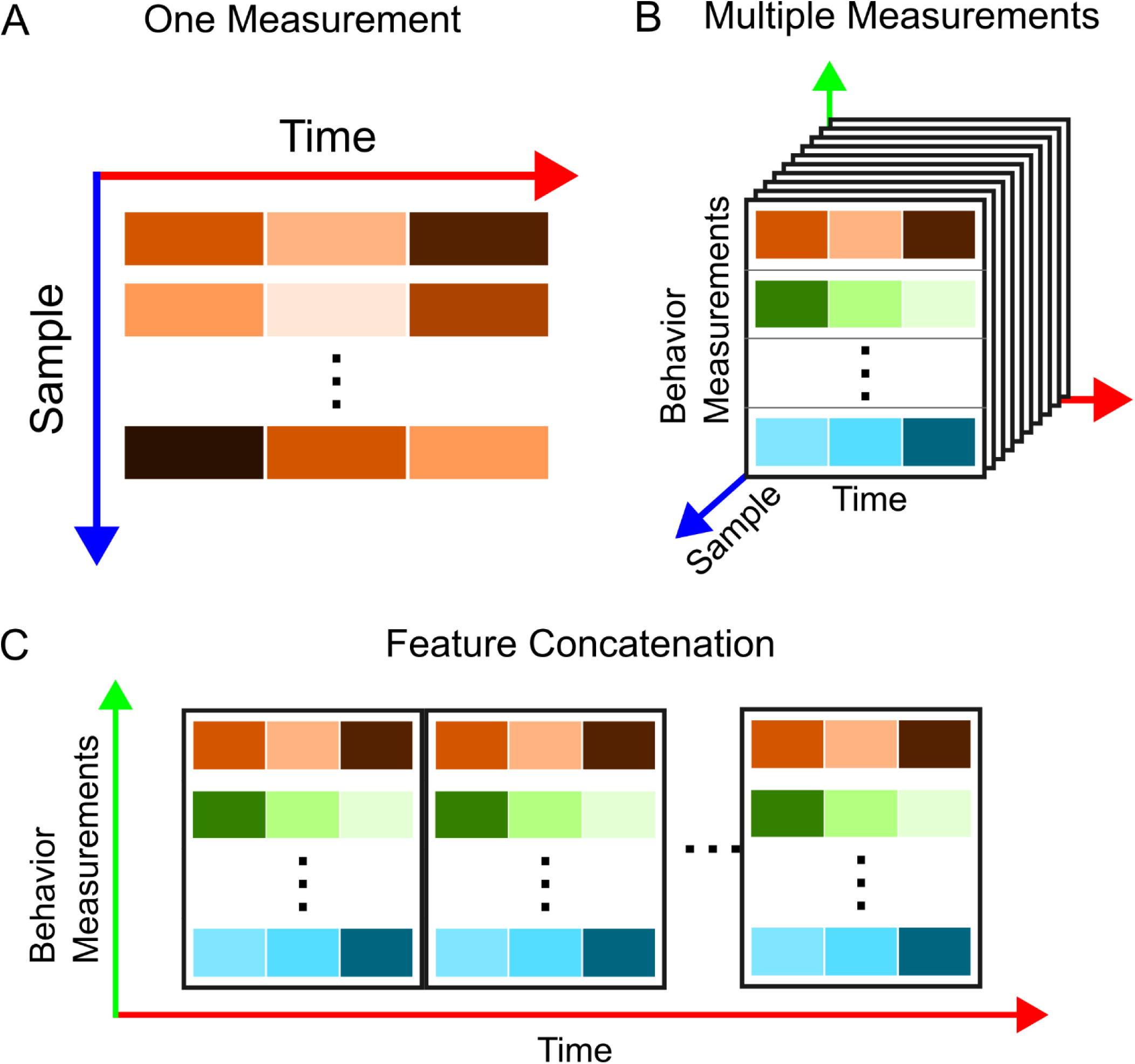
Schematic structures of behavior data in the forms of multivariate data and the multi- dimensional tensor. **(A)** The schematic structure of a time-series behavior data with one single behavior measurement repeatedly collected from multiple samples (blue axis) over time (red axis). The resulting data structure is a matrix. Each brown rectangle represents a unique measurement per sample per time point. Its color intensity represents the magnitude of the measurement. **(B)** The schematic structure of a time-series behavior data with multiple behavior measurements (green axis) repeatedly collected from multiple samples (blue axis) over time (red axis). The resulting data structure is a three-dimensional array known as a three-dimensional tensor (3DT) or a third-order tensor, a type of multi-dimensional tensor (MDT). In this array, each “sheet” represents the data collected from one sample. The different color rectangles (brown, green, and blue) represent different behavior measurements per sample per time point, whereas their color intensities represent the magnitude of the measurement. **(C)** The schematic structure of MDT data when matricized by Feature Concatenation. The matricization process concatenates features by joining each “sheet” or each slice of the 3DT shown in (B). Here, the sample dimension is reduced along the time dimension (red axis), transforming the 3DT into a two-dimensional tensor or a matrix.

When this happens, this data structure can be generalized into a multi-dimensional tensor (MDT), also defined as multi-order tensors in tensor terminology [13–15] (see Multi-Dimensional Tensor section below). In this study, we refer to MDT as tensors with more than two dimensions. Since the MDT structure contains more than two dimensions, it is incompatible with MVA. For example, MANOVA cannot directly use the MDT data as input. Hence, the MDT structure poses a challenge to data analysis in behavior studies.

This challenge is partially resolved by transforming the MDT to a two-dimensional tensor (2DT) and reducing the number of dimensions. This transformation makes the transformed data structure compatible with MVA. The tensor dimensions can be reduced by many models, such as tensor component analysis (TCA) based on CANDECOMP/PARFAC (CP) tensor decomposition [13,16,17]. These models are generalizations of 2DT data dimensionality reduction. For instance, TCA is conceptually similar to the principal component analysis (PCA) [17]. However, these models have not been widely applied to analyzing animal behaviors, possibly due to the difficulties in interpreting their mathematical representations, operations, and resulting outputs. The same interpretability issue occurs in PCA, where the resulting PCs are the mathematical representation of multiple original features. The abstract representation of the features makes it hard to interpret the significance and effect of the original features [18–20]. To mitigate this interpretability issue, many behavioral scientists reduce the tensor dimensions in their data with alternative approaches. One approach called matricization transforms features in MDT space into new features in a less complex 2DT space [13,21], which makes the resulting features more interpretable and compatible with MVA. There are two matricization strategies commonly used in behavior studies, and we refer to them as 1) Index Construction and 2) Feature Concatenation. In Index Construction, a single index is constructed to represent all or a subset of the original features in the MDT [22]. For example, the average of a group of animal displacement can be calculated to reduce the sample dimension, assuming the samples in the group are homogenous [23–25]. In Feature Concatenation, all or a subset of features in a dimension of MDT dimension are selected to form simpler 2DTs or matrices (e.g., the individual sheets in Fig 1B) [26]. These matrices are then joined or concatenated, essentially flattening the complex data “cube” (Fig 1B) into a long data “sheet” (Fig 1C) [13,21,27] (See the next section for the technical details). For example, neural recordings across different trials can be joined together and flatten the 3DT neural recording data onto other dimensions [17,28]. Once the MDT data are reduced to 2DT data, the data can be analyzed by MVA. However, both matricization strategies have drawbacks. For example, Index Construction may exclude information from the original features that may be essential to describe the behavior dynamics, whereas Feature Concatenation may create too many transformed features and introduce additional noise to the downstream analysis. These issues may impact the performance of downstream MVA. Hence, both matricization strategies and their transformed features must be evaluated to select the best strategy and set of features that optimize the MVA performance.

In this study, we applied several matricization approaches to analyze an MDT dataset obtained from a zebrafish behavior assay. We empirically evaluated how different matricization and feature-selection strategies may affect the performance of MVA, specifically for classification. We found that the 2DT features selected by Feature Concatenation improved the classification performance the most. In the following sections, we provide the relevant theoretical background on MDT and feature selection.

### Multi-dimensional tensor

In this study, an MDT is defined as a tensor with higher than two dimensions. For instance, a 3-dimensional tensor (3DT) is represented in Fig 1B and can be denoted as ℝ^*N×M×S*^. In animal-behavior studies, *N* represents the number of samples, *M* represents the number of behavior measurement types, and *S* represents the number of time points. In other words, this 3DT can be considered a higher-order matrix or a matrix of matrices [13]. The 3DT can be transformed into a 2DT or a matrix using matricization methods such as Index Construction or Feature Concatenation. For example, to reduce the behavior-measurement dimension by Index Construction, a behavior index is constructed to represent all or a subset of *M* features, reducing the number of features from *M* to 1. The resulting 2DT is ℝ^*N×S*^ denoting the transformed dataset, equivalent to a matrix of *N* rows by *S* columns (as illustrated in Fig 1A). To reduce the behavior-measurement dimension by Feature Concatenation, the data is sliced by the number of features in a dimension to generate sub-matrices which are then joined to create one matrix. For example, in the case of Fig 1C, the concatenation essentially flattens the 3DT in Fig 1B to a 2DT with the transformed structure of ℝ^*M×NS*^, equivalent to a matrix of *M* rows by *N* × *S* columns. To reduce the time dimension the time series can be transformed into symbolic representations [29,30] or behavior motifs [31,32]. In this case, the resulting matrix may contain *N* rows by *M* × *L* columns, where *L* denotes the reduced number of time points, i.e., *L* < *S*.

### Feature selection

After the matricization of an MDT, the transformed features in the 2DT space must be assessed to determine which ones effectively capture the behavioral dynamics. Only the informative features should be used downstream MVA to optimize the analysis performance. The informative features can be selected by 1) filter method, 2) embedded method, and 3) wrapper method [33,34] (Table 1). The choice of method often relies on empirical experience and practical constraints [35].

**Table 1.** Three Methods for Feature Selection.

| Method | Approach | Examples | Advantages | Disadvantages |
| --- | --- | --- | --- | --- |
| Filter | The filter method uses quantified metrics to rank and filter features without a machine-learning process [33,34,36]. | P-value by statistical testing [37,38]<br>Fold change [38]<br>Correlation [39,40]<br>Information Gain [33,41] | Low computational cost [33,34]<br>Independent from the machine-learning processes [33,34] | Neglects dependencies among features [33,34] |
| Embedded | The embedded method relies on a built-in mechanism in a learning machine where learning and feature assessment are intertwined [33,34]. | LASSO [42]<br>Random Forest [43]<br>Decision Tree [44] | Relatively low computation cost [33,34]<br>Consider dependencies among features [33,34] | Feature importance depends on the mechanism and limitation of the classifier [33,34] |
| Wrapper | The wrapper method evaluates feature importance by its performance in a learning machine [33,34,36]. | Sequential Forward Selection [45]<br>Sequential Backward Elimination [45] | Consider dependencies among features [33,34]<br>Superior performance for supervised machine learning [33,35,46] | High computation time for exhaustive search [33,34]<br>Feature importance depends on the mechanism and limitation of the classifier [33,34] |

To harness the power of all three methods and mitigate their drawbacks, some feature-selection workflows combine features selected by these methods through intersection or union operations, an approach known as the ensemble approach [47,48]. The ensemble approach via intersection has been shown to select features from complex medical datasets that achieve the best classification accuracy in empirical testing [48]. Therefore, it is possible that the ensemble approach can also be applied to complex behavior datasets. In this study, we demonstrated the feasibility of this approach by matricizing along the behavioral-measurement dimension. The wrapper method in the ensemble approach may be less compatible with behavior studies because the recordings may take hours or days with a time resolution in seconds or milliseconds. There are too many time points for the wrapper methods, especially neural networks, to iterate exhaustively through under limited computational resources.

Therefore, we evaluated an ensemble framework that utilizes filter and embedded methods after reducing the dimensionality of an MDT. In our ensemble approach, the 2DT features were aggregated by intersection and union operations. The performance of both operations was evaluated by a classification task to analyze zebrafish visual behavior.

## Methods

### Zebrafish husbandry and breeding

The zebrafish used in this study were the AB strain obtained from the Zebrafish International Resource Center <https://zfin.org/ZDB-GENO-960809-7>. The visually-impaired transgenic line, *Tg(rho:Hsa.RH1_Q344X)*, was generated and used in our previous studies and is referred to in this study as Q344X [23,49]. The animals were bred and maintained following standard procedures [50,51]. Prior to embryo collection, adults were sex-separated in breed tanks the night before breeding. They were mixed and bred from 9 a.m. to 11 a.m. the following day. All collected embryos were raised in the E3 medium at 28 °C in an incubator with the same light-dark cycle (14-hour light/ 10-hour dark) as in the fish facility. The medium was changed every day. During the process, unhealthy and dead embryos were discarded. All protocols were approved by the Purdue University Institutional Animal Care and Use Committee.

### Visual-motor response assay

The MDT dataset used in this study was generated by the zebrafish visual-motor response (VMR) assay [52,53], a systematic approach to measure the visual startle (i.e., swimming behavior) of zebrafish larvae triggered by drastic light changes. In the VMR assay, the larvae are individually placed in a 96-well plate, and their visual startles are recorded by an infrared camera in a Zebrabox System (Viewpoint Life Sciences). Prior to the VMR assay, zebrafish larvae at 6 days postfertilization (dpf) were transferred to the E3 medium in Whatman UNIPlate square 96-well plates (VWR). The 96-well plates with larvae were placed in light-proof boxes for 14 hours of dark habituation inside the incubator. At 7 dpf, these 96-well plates were placed inside the Zebrabox for data collection. The larval VMR was triggered by two types of light stimulation: an onset (light-on) and an offset (light-off). The videos were recorded in the tracking mode, binning the behavioral measurements every second. For the light-on VMR, the protocol consisted of a 1-hour dark period and a 5-minute (300-second) light period. For the light-off VMR, the protocol consisted of a 1-hour dark period, a 1-hour light period, and a 5-minute (300-second) dark period. The VMR data used in this study were collected during our ongoing drug screening for retinitis pigmentosa using the Q344X mutant [23,49] to detect drug improvement in rod response [54]. The scotopic light intensity was fine-tuned to detect the maximal difference in the light-on and light-off VMR between WT and the Q344X mutant. The total irradiance for light stimulation over the visible spectrum was 0.00965 µW/cm^2^ and 0.000389 µW/cm^2^ for the light-on and light-off VMR, respectively. All experiments were conducted between 9 a.m. and 6 p.m. to minimize the effect of circadian rhythm on vision [23,52].

### VMR data processing and analysis

#### General

All data analyses in this section were conducted with R statistical environment version 4.1.2 < https://www.r-project.org/>, except for data extraction from the VMR videos, which was conducted with ZebraLab version 3,22,3,77, and the multilayer perceptron classifier, which was conducted with Python version 3.9 <https://www.python.org/>. All the raw VMR data used in this study can be accessed through the Harvard Dataverse via the following URL: <https://doi.org/10.7910/DVN/B8HBU9>. The code for the analysis can be accessed through GitHub via the following URL: <https://github.com/wang4537/VMR-Feature-Selection>.

### VMR data structure

The larval activity in the recorded video was extracted and quantified into nine behavioral measurements (Fig 2A). First, the larval movement was categorized into three activity levels: inactivity (*ina*; speed < 0.6 cm/s), small activity (*sml*; speed ≥ 0.6 cm/s and < 1 cm/s), and large activity (*lar*; speed ≥ 1 cm/s). Speed was calculated based on every 5 frames of the video recording using the following equation, according to the specifications of Viewpoint Zebralab:

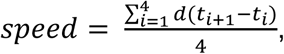

where *t_i_* denotes the location of the animal center mass at Frame *i*; *d(t_i+1_-t_i_)* denotes the distance of the animal center mass between Frame *t_i+1_* and Frame *t_i_*. Second, within each activity category, three measurements were collected: duration of movement (*dur*), count of speed change (*ct*), and distance of movement (*dist*). Hence, nine behavioral measurements were taken per larva per second (Fig 2A). The complete definitions of these nine measurements are listed in the S1 Table.

**Fig 2.**
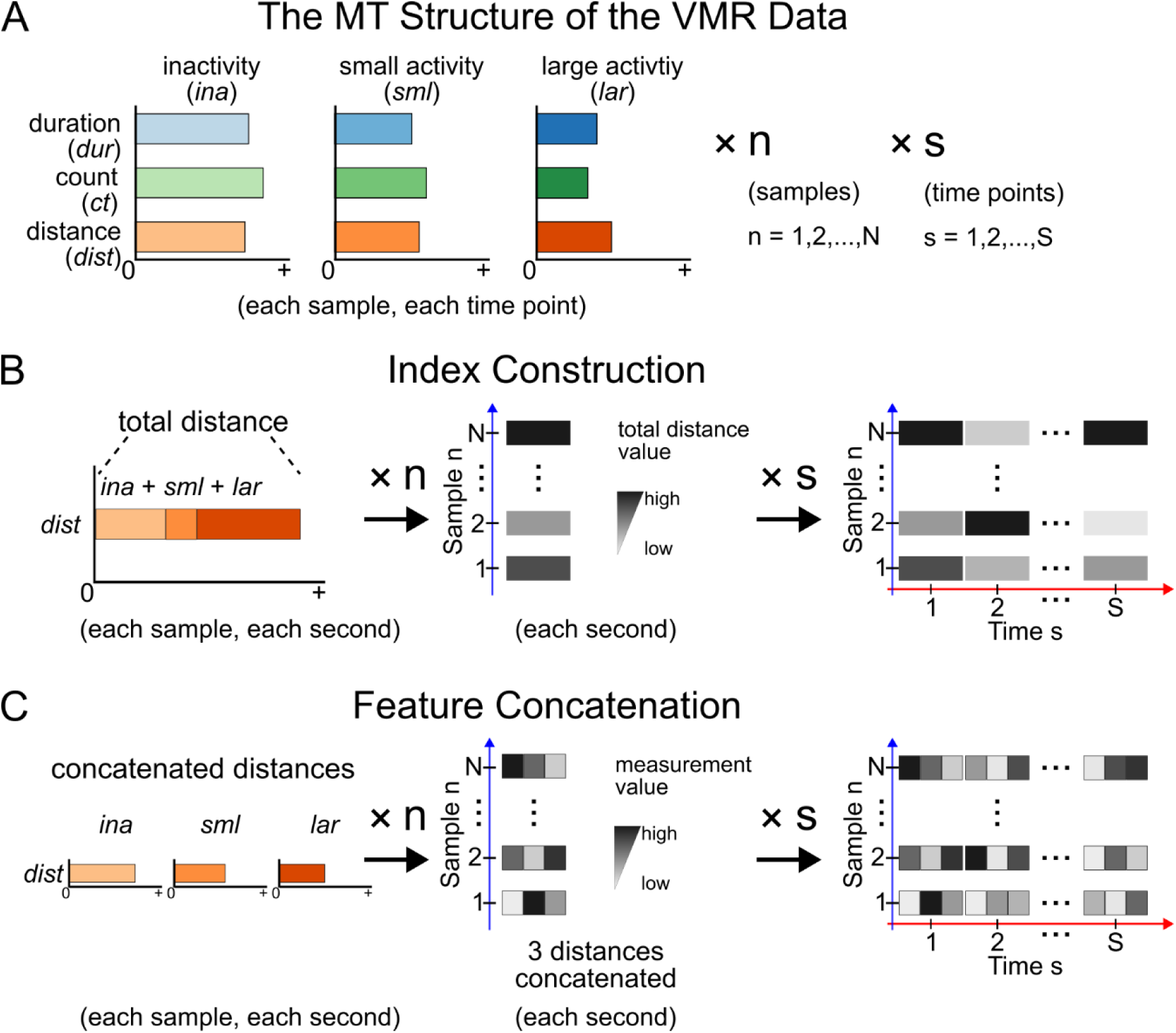
The schematics of the 3DT from the zebrafish behavior data used in this study and its matricization strategies. **(A)** The schematic of the 3DT of the VMR data. In the VMR assay, zebrafish behaviors were recorded by nine behavior measurements. Three types of behavior were recorded: duration (*dur)*, count (*ct)*, and distance (*dist*), each at three activity levels: inactivity (*ina;* < 0.6 cm/s), small activity (*sml;* ≥ 0.6 cm/s and < 1 cm/s), and large activity (*lar;* ≥ 1 cm/s). Each behavior measurement is represented by a unique color, whereas the activity levels of that behavior are represented by different color intensities. The size of the bars indicates the magnitude of the activities. See Table S1 for the detailed definitions. These nine measurements were collected from each sample at each second. In a time-series experiment (s) with multiple samples and treatment conditions (n), this collection scheme results in 3DT data. **(B)** The schematic of Index Construction by a Total Distance index. This index is the sum of the *dist* measurements at three activity levels, i.e., *ina, sml,* and *lar*, for each sample at each second (*left*). When this index is calculated for all samples (middle) over time (right), this results in a simpler data structure with one Total Distance index instead of nine behavior measurements. In the middle and right panels, each bar represents a Total Distance index for a sample in a second, whereas the darkness of the grey tone represents the magnitude of the index. **(C)** The schematic of Feature Concatenation by concatenating *dist* measurements. The left panel represents the *inadist, smldist,* and *lardist* from the original data (A). These features were joined/concatenated at each second. When this is performed for all samples (middle) over time (right), this results in a simpler data structure with data arranged in a 2DT space rather than the original 3T space (A). In the middle and right panels, each bar represents the three concatenated features in three grey tones for a sample in a second, whereas the darkness of the grey tone represents the magnitude of the behavior measurement. Prior to the matricization, the VMR data was first normalized using the established VMR method (See S1 Appendix for details).

### Matricizing the 3DT VMR data

Before matricization, the VMR data were first normalized to remove confounding factors, including light intensity variations, batch effect, and baseline activity variations [55] (See S1 Appendix for details). Then, the normalized 3DT VMR data were matricized by Index Construction and Feature Concatenation. In Index Construction, an index was calculated to represent part or all of the behavior measurements. For instance, a commonly used index in the VMR data is the Total Distance [23,56–58], the sum of distance (*dist*) across all three activity levels (i.e., *ina*, *sml*, and *lar*) for each sample (or larva) at each second for all time points (Fig 2B). In other words, the Total Distance is calculated as follows:

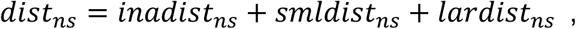

where *dist_ns_* denotes the Total Distance for sample *n* at time point *s*, *n=1,2,…,N* and *s=1,2,…,S*. Here *N* denotes the total number of samples, *S* denotes the total number of time points, and *inadist_ns_*, *smldist_ns_*, and *lardist_ns_* denote the inactivity, small activity, and large activity distance, respectively, for sample *n* at time point *s*. Calculating the *dist* behavior index reduces the number of behavior measurements from 9 to 1 by merging 3 *dist* measurements and discarding the remaining 6 measurements. This operation transforms the 3DT to a 2DT of *N×S* dimensions with *S* features. In this study, we named this type of 2DT feature with *distance* and the time point after the light change, separated by a dot (“.”). For example, *distance.1* refers to the total-distance feature at 1 second after light change for each larva. We also tested two other indices, L1 Norm and L2 Norm, which utilize all 9 features instead of discarding some features that may contain unique information. In data science, L1 and L2 Norms are used to reduce the dimensionality of the multivariate data [59]. Mathematically, L1 and L2 Norms are defined as follows:

Given a vector:

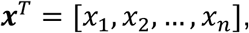

where *x_1_*, *x_2_*, …, *x_n_* represents values for each dimension. The L1 Norm of ***x*** is defined as:

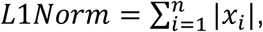

and the L2 Norm of ***x*** is defined as:

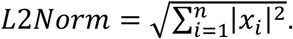

In our VMR data, the *n*th larval movement at *s*th second can be regarded as a vector of size *M*. The corresponding L1 and L2 Norms can be calculated as follows:

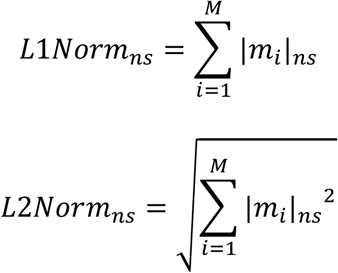

where *m_i_* denotes the *i*th behavior measurements, where *i = 1,2,…M*; *n* denotes the *n*th sample, where *n=1,2,…,N*; *s* denotes the *s*th time point, where *s=1,2,…,S*. Since behavior measurements were in different units and scales, the values were first standardized to their corresponding mean and standard deviation. After standardization, L1 and L2 Norms were calculated as the 2DT features. The 2DT features were named following the same convention used in the Total Distance index. For example, *L1Norm.1* refers to the L1 Norm feature at 1 second after light change for each larva.

In Feature Concatenation, a few or all behavior measurements were concatenated per time point per sample to reduce the behavior measurement dimension, as illustrated in Fig 1C. In this study, we reduced the behavior-measurement dimension by two Feature Concatenation approaches: Concatenated 3 Distances (C3D) and Concatenated 9 Measurements (C9M). In C3D, we joined all distance-related measurements (i.e., *inadist*, *smldist*, and *lardist*) for each larva at each second (Fig 2C). After the join of each *dist* measurement, the transformed matrix will have the dimension of *N×(3×S)* with *3×S* features in the 2DT space, where *N* denotes the size for the sample dimension and *S* denotes the size for the time dimension. In C9M, we joined all features in the behavioral-measurement dimension. After the join of all measurements, the transformed matrix will have the dimension of *N×(9×S)* with *9×S* features in the 2DT space. In both 2DT matricized by either C3D or C9M, each 2DT feature was named using the behavior measurement and the time point after the light change, separated by a dot (“.”). For example, *inadist.1* refers to the feature representing the *inadist* (inactivity distance) at 1 second after light change for each larva.

### Feature selection

The 2DT features were selected by aggregating the results of filter and embedded methods. For the filter method, two metrics were used: the p-value of the Wilcoxon rank sum test [60,61] and the average fold change (FC) [62]. The p-value was calculated by running a Wilcoxon rank sum test between the WT and Q344X larvae using each feature and adjusted for multiple hypothesis testing using Holm’s method [63]. A feature was deemed important and selected if the adjusted p-value was less than 0.01. The FC were calculated for each feature by the following formula:

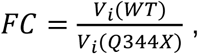

where *V_i_(WT)* denotes the average value of feature *i* in all WT larvae; *V_i_(Q344X)* denotes the average value of feature *i* in all Q344X larvae; *i = 1, 2, …, S* for features matricized by Index Construction, and *i = 1, 2, …, M*×*S* for features matricized by Feature Concatenation. For both Index Construction and Feature Concatenation, *S* denotes the number of time points; *M* denotes the number of behavior-measurement types. A feature was selected if the absolute of its log2(FC) was greater than 0.38 in light-on VMR (equivalent to a linear FC > 1.3 or < 0.77 [i.e. 1/1.3]) or 0.58 in light-off VMR (equivalent to a linear FC > 1.5 or < 0.67 [i.e. 1/1.5]).

These cutoffs were empirically chosen to balance the stringency of selection and the number of selected features based on our previous observations that the WT larvae typically display a lower scotopic VMR under the light-on stimulus [54]. For the embedded method, Random Forest (RF) [43] was used. RF ranked the feature importance by the Mean Decrease of the Gini Index (MeanDecreaseGini) of each feature [64,65]. MeanDecreaseGini is calculated when RF randomly splits all features into random subsets and assesses information impurity for the features [64,66]. Finally, the features selected by the filter and embedded methods were aggregated by either intersection or union operation to create intersection and union sets.

### Evaluation of feature selection and matricization performance

The performance of the different selected feature sets and matricization methods was evaluated by a classification task to distinguish the WT and the Q344X mutant. The following classifiers were used: Support Vector Machine (SVM), Naïve Bayes (NB), K-Nearest Neighbor (KNN), Decision Tree (DT), RF, Extreme Gradient Boosting (XGB), and Multilayer Perceptron (MLP). SVM, KNN, and NB are classic classifiers previously used for VMR analysis [67]. Tree-based methods, including DT, RF, and XGB, are considered superior in performance to the classic classifiers in other fields [68,69]. MLP is a type of neural-network classifier [70] that is adaptive and capable of non-linear relationships in the data, achieving high accuracy and robustness [71]. Detailed information on these packages and their hyperparameter selection can be found in S2 Table. For each classifier, five feature sets were used: Full set, features selected by the filter method only (Filter), features selected by the embedded method only (Embedded), intersection of the features selected by the Filter and Embedded methods (Intersect), and union of the features selected by the Filter and Embedded methods (Union). The light-on and light-off VMR were analyzed separately.

The performance of the classifiers on each feature set was evaluated using a holdout validation (HV) approach with 10-fold cross-validation (CV) tuning (Fig 3). First, 80% of the dataset was allocated as the training set. This training set was used in the 10-fold CV to tune each classifier’s hyperparameters and assess the performance of each feature set under different matricization methods. A range of hyperparameters per classifier was iterated using a grid search method [32,38], with the area under the receiver operating characteristic (AUROC) curve as the performance metric. The best hyperparameters for each classifier and feature set were selected based on the highest average AUROC score obtained in the CV. Next, the remaining 20% of the data (the testing set) was used in the HV to evaluate the generalizability of the feature sets and matricization methods. Each classifier was trained using the training set with the hyperparameters identified as optimal in the CV and then applied to the testing set.

**Fig 3.**
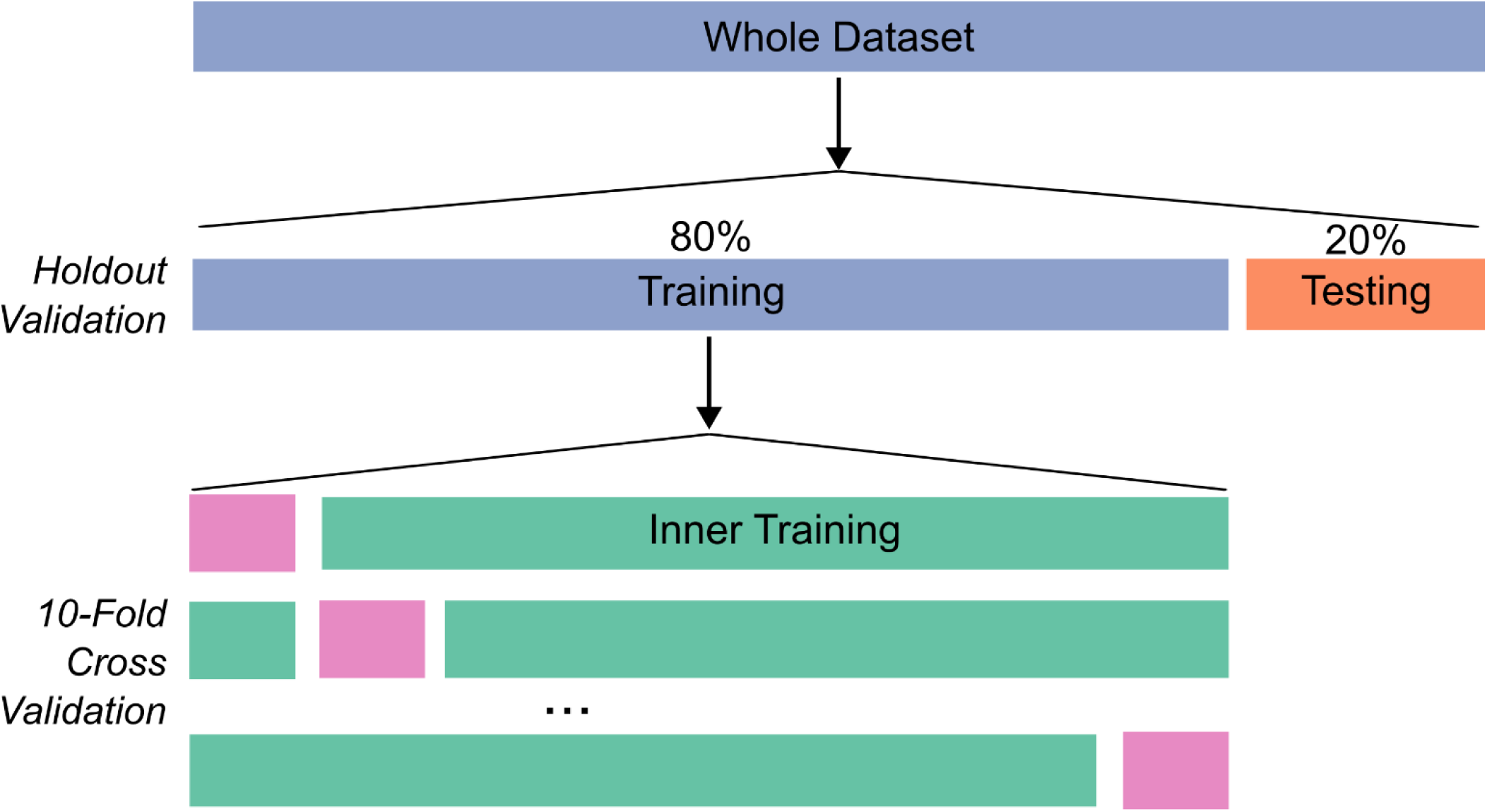
Evaluation of different matricization and feature-selection methods. The schematic diagram shows the design of the performance evaluation of different feature-selection strategies by a classification task using the VMR data. Two evaluations were conducted: holdout validation (HV) and 10-fold cross-validation (CV). In HV, the whole dataset was split into two subsets: the training set (80% of the data, blue) and the testing set (20% of the data, orange). The training set was used to select features by various methods described in the text and as input for the CV. In CV, the training set was split to tune hyperparameters and compare performances of different matricization methods and feature sets. Each fold consists of inner training data (cyan) to train the classifier and then tested on the validation set (magenta), evaluated by area under receiver operating curve (AUROC). The AUROC across all folds were then averaged. The highest average AUROC across the hyperparameter range was used to determine the optimal hyperparameters and represent the performance per feature set per matricization method in CV. Then, the optimal hyperparameters were utilized to build classifiers using the entire training data (blue) to test performances on the testing set (orange) in HV to validate the generalizability of the matricization and feature-selection methods.

Classification performance on the testing set was measured using accuracy, sensitivity, specificity, precision, and the Kappa statistic [69,70].

## Results

### The 3DT VMR data and classic dimension reduction

The 3DT behavior data used in this study consisted of a light-on and a light-off VMR dataset, each with 9 biological replications of the VMR collected from 24 WT and 24 Q344X larvae at 7 dpf. In the VMR protocol, the final session was a 5-minute (300- second) light-on or light-off period for triggering the light-on and light-off VMR, respectively. Hence, the analysis focused on this 5-minute period. The data structure followed the one outlined in Fig 2A and has 9 behavior measurements (*M = 9*), 432 samples (*N = 432*), and 300 seconds after light change (*S = 300*). Therefore, our input MDT has dimensions of *432×9×300.* This data was matricized to 2DT data by Index Construction or Feature Concatenation. For instance, when a Total Distance index was constructed (Fig 2B), 3 distance measurements were reduced to 1 distance index, while the 6 other measurements were discarded. This operation transformed the VMR data from 3DT to 2DT format of *432×300* dimensions. The Total Distance is a commonly used index for this type of data, as it provides an intuitive way to visualize the behavioral and biological differences between sample types [23,56–58]. For example, the visually-impaired Q344X larvae lacked the visual startle immediately after light onset (Fig 4A) and light offset (Fig 4B), compared to the WT larvae. They also displayed a lower sustained response after light offset (Fig 4B). These intuitive differences allow for setting up simple rules and statistical tests to detect differences between genotypes, including using the fast response in the first few seconds to study visual startle [8,23,54]. In addition to Total Distance, the data can also be matricized by concatenating the 3 distance measurements along the time tensor, transforming the data to a matrix of *432×(3×300)* or *432×900* in size (Fig 2C). This matrix can also be used for downstream MVA.

**Fig 4.**
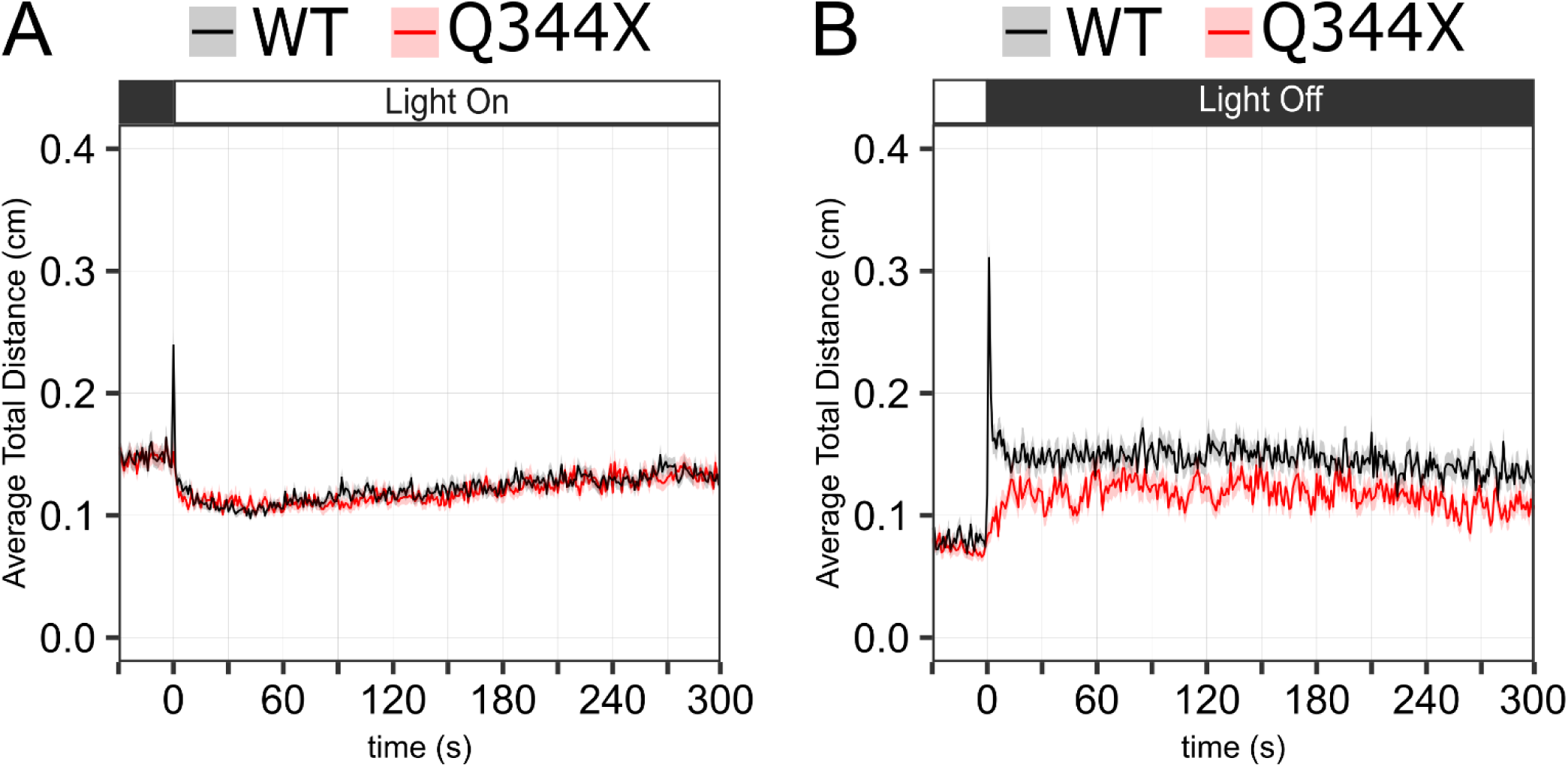
The light-on and light-off VMR of the WT and Q344X larvae as illustrated by the Total Distance index. **(A)** The light-on VMR of the WT (black trace) and Q344X (red trace) larvae. Each trace represents data collected from 9 biological replicates with 24 larvae per genotype per replicate. The plot shows the average Total Distance traveled per second by the two genotypes from 30 seconds before light onset at 0 to 300 seconds after light onset. The color ribbon on each trace represents the standard error of the mean (SEM). **(B)** The light-off VMR of the WT (black trace) and Q344X (red trace) larvae. Each trace represents data collected from 9 biological replicates with 24 larvae per genotype per replicate. The plot shows the average Total Distance traveled per second by the two genotypes from 30 seconds before light offset at 0 to 300 seconds after light offset. The color ribbon on each trace represents the SEM. The black and white boxes on each plot represent the light-off and light-on period.

In contrast, the Tensor Component Analysis (TCA) extracted the differences between data points along each dimension, but the results were less intuitive (S1 Fig). For example, it revealed several unique patterns in each dimension, but they do not necessarily differentiate or pinpoint the VMR difference between the WT and Q344X larvae. Hence, TCA results are harder to interpret when compared to matricization, especially Index Construction. On the contrary, matricization can give rise to features that are easier to interpret. However, these features may not always carry relevant information for downstream analysis. Hence, their importance was further evaluated by feature selection with the filter and embedded methods.

### Feature selection—filter method

The filter method was used to select 2DT features from the light-on and light-off VMR datasets matricized by Index Construction (Fig 5) and Feature Concatenation (Fig 6). Fig 5 shows the results of Index Construction by three indices: Total Distance, L1 Norm, and L2 Norm. In the light-on VMR dataset, the filter method only selected *distance.1* (i.e., the Total Distance in the first second) out of the 300 features from the Total Distance index (Fig 5A). It did not select any features from L1 or L2 Norm indices (Fig 5B & C). In the light-off VMR dataset, the filter method selected 26 out of the 300 features from the Total Distance index (Fig 5D). Among these 26 features, *distance.2* and *distance.3* had the highest *–log_10_(p)* and *log_2_(FC)* values. The filter method did not select any feature from L1 Norm index, but it selected 3 out of 300 features from the L2 Norm index, namely *L2Norm.2*, *L2Norm.3*, and *L2Norm.7*.

**Fig 5.**
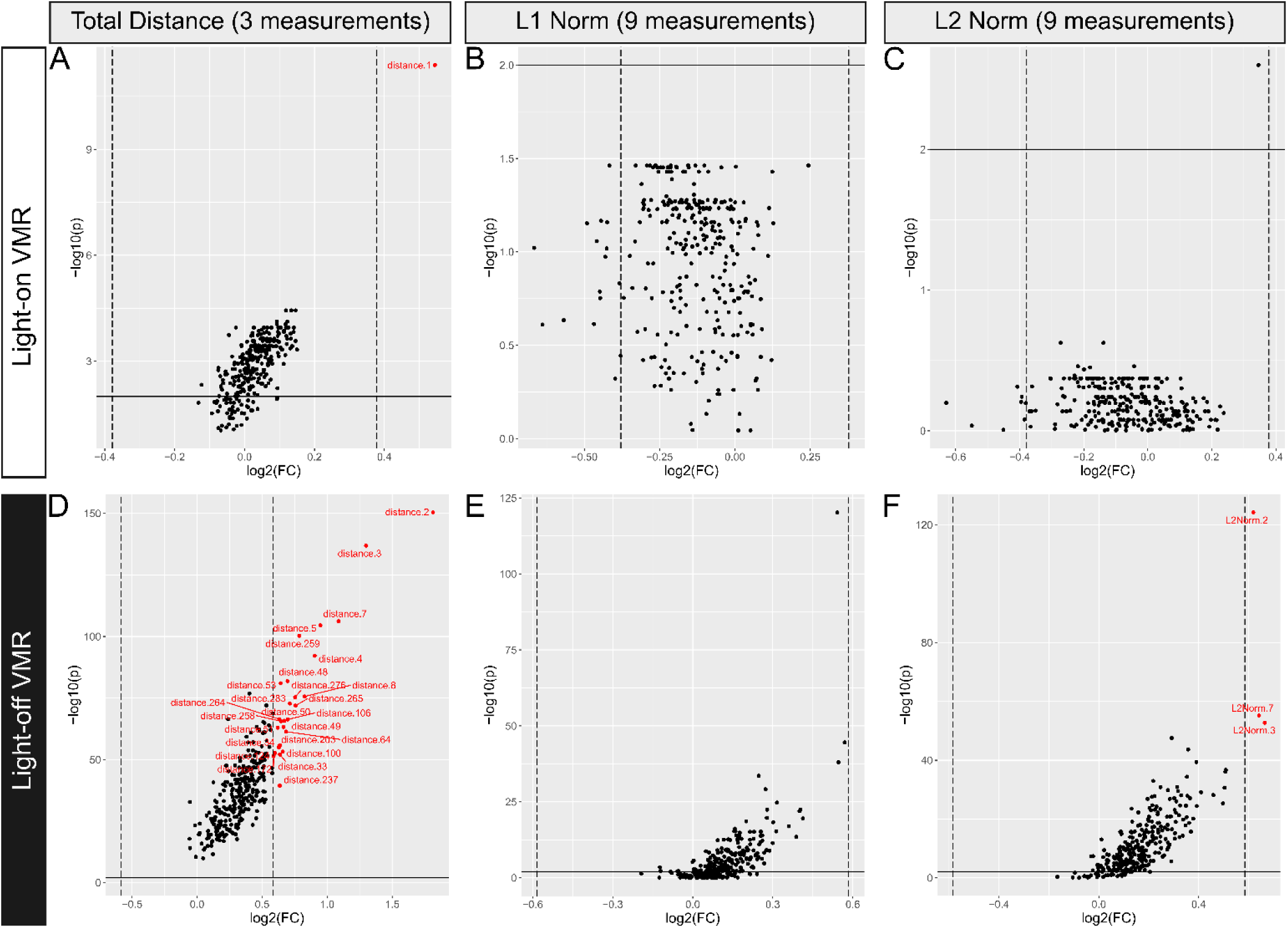
Using the filter method to select features from the VMR datasets matricized by Index Construction. (A–C) Volcano plots showing the features from the light-on dataset matricized by the Total Distance, L1 Norm, and L2 Norm indices, respectively. **(D–F)** Volcano plots showing the features from the light-off dataset matricized by the Total Distance, L1 Norm, and L2 Norm indices, respectively. A feature was deemed significant and selected from the matricized indices if the Holm adjusted p-value from the Wilcoxon rank sum test (in -log_10_(p); Y-axis) and the fold change (in log_2_(FC); X-axis) of that feature between the WT and Q344X mutants exceed the cutoffs. The cutoff for p-value is -log_10_(p) > 2 (i.e., p < 0.01) for both the light-on and light-off VMR (horizontal solid lines), whereas the cutoffs for FC were |log2(FC)| > 0.38 and |log2(FC)| > 0.58 for the light-on and light-off VMR, respectively (vertical dash lines). The selected features were highlighted by red dots if the WT’s values were significantly larger than the values of the Q344X mutant. They were highlighted by black dots, if the feature values between the WT and the Q344X were not significantly different. The feature label indicates the name of the index and the corresponding time point after the light change, separated by a dot (“.”). For example, the “*distance.1*” refers to the Total Distance in the first second after the light change.

**Fig 6.**
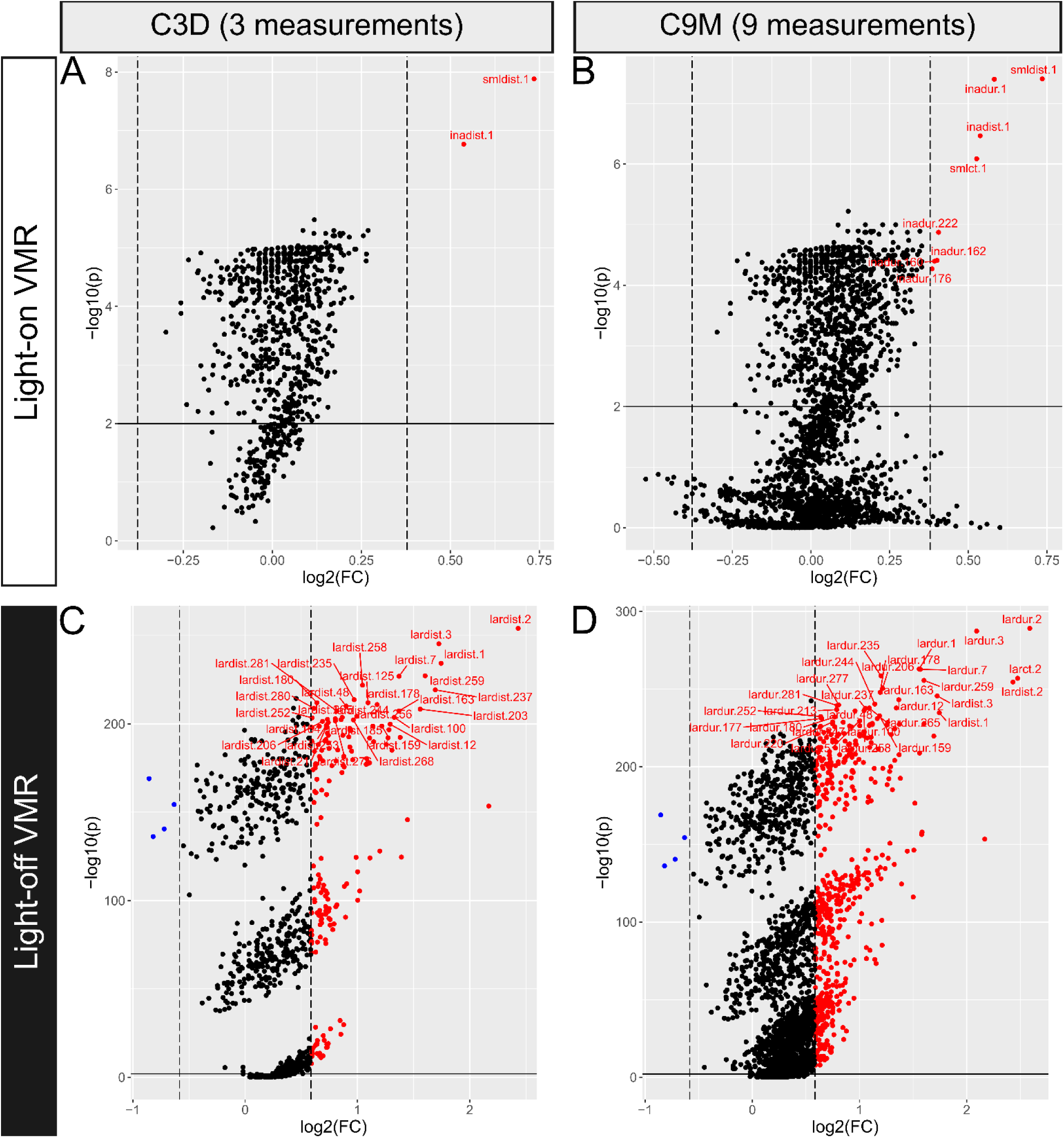
Using the filter method to select features from the VMR datasets matricized by Feature Concatenation. (A–B) Volcano plots showing features from the light-on VMR dataset matricized by C3D and C9M, respectively. **(C–D)** Volcano plots showing features selected from the light-off VMR dataset matricized by C3D and C9M, respectively. The features were selected using the same filter method and cutoffs as in Fig 5. They were also labeled in the same style as in Fig. 5, except for the blue dots, which highlighted selected features where the WT’s values were significantly lower than those of the Q344X mutant.

Fig 6 shows the feature-selection results of Feature Concatenation by C3D and C9M. In the light-on VMR data, the filter method selected 2 features, *smldist.1* and *inadist.1*, out of the 900 features (3 measurements x 300 s) from C3D (Fig 6A). It also selected 8 out of the 2700 features (9 measurements x 300 s) from C9M (Fig 6B). Among the selected 9 features, *smldist.1*, *inadur.1*, and *inadist.1* had the highest *–*log_10_(p) and |log_2_(FC)| values. In the light-off VMR, the filter method selected 179 out of 900 features from C3D (Fig 6C). Among these 179 features, *lardist.2*, *lardist.3* and *lardist.1* had the highest –log_10_(p) and log_2_(FC) values. Four selected features had negative log_2_(FC) values (blue dots, Fig 6C), including *lardist.41*, *lardiest*.*92*, *lardiest*.*183,* and *lardist*.*233*, indicating that the feature values were higher in the Q344X mutant larvae than in the WT larvae. The filter method also selected 548 out of 2700 features from C9M (Fig 6D). Among these 548 features, *lardur.2*, *lardur.3*, *larct.2,* and *lardur.1* had the highest –log_10_(p) and |log_2_(FC)| values. Four selected features also had negative log_2_(FC) values (blue dots, Fig 6D), and they are the same 4 features selected from C3D in Fig 6C.

### Feature selection—embedded method

Next, the embedded method was used to select 2DT features from the light-on and light-off VMR datasets matricized by Index Construction (Fig 7) and Feature Concatenation (Fig 8). Fig 7 shows the results of Index Construction by three indices: Total Distance, L1 Norm, and L2 Norm. In the light-on VMR dataset, the embedded method selected *distance.1* and *distance.189* out of the 300 features from the Total Distance Index (Fig 7A). The method also selected *L1Norm.1*, *L1Norm30*, and *L1Norm.205* out of the 300 features from the L1 Norm index (Fig 7B), and *L2Norm.1* and *L2Norm.45* out of the 300 features from the L2 Norm index (Fig 7C). In the light-off VMR dataset, the embedded method selected 8 out of 300 features from the Total Distance index (Fig 7D). Among these 8 features, *distance.2*, *distance.3*, and *distance.5* had the highest MeanDecreaseGini values. The embedded method also selected 6 out of the 300 features from the L1 Norm index (Fig 7E). Among these 6 features, *L1Norm.2*, *L1Norm.3,* and *L1Norm.7* had the highest values of MeanDecreaseGini. In addition, the embedded method selected 6 out of the 300 features from the L2 Norm index (Fig 7F). Among these 6 features, *L2Norm.2*, *L2Norm.3,* and *L2Norm.7* had the highest MeanDecreaseGini values.

**Fig 7.**
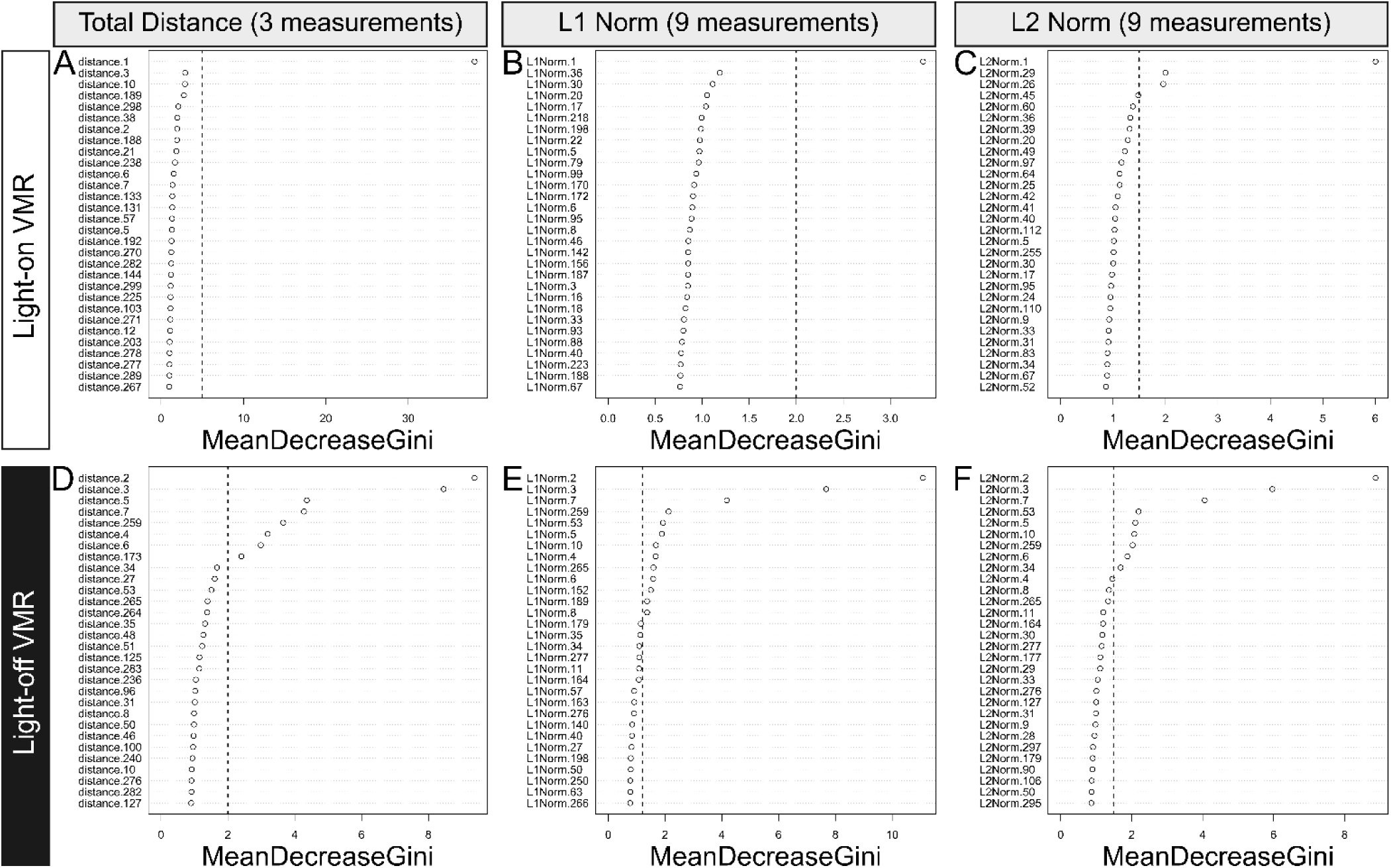
Using the embedded method to select features from the VMR dataset matricized by index construction. (A–C) Top 30 features from the light-on VMR dataset matricized by the Total Distance, L1 Norm, and L2 Norm indices. **(D–F)** Top 30 features from the light-off VMR dataset matricized by the Total Distance, L1 Norm, and L2 Norm. These features (Y-axis) were named as in Fig. 5 and ranked by Random Forest using the mean decrease of the Gini index (MeanDecreaseGini; X-axis) in descending order. In each panel, a feature was selected if its MeanDecreaseGini value was larger than the cutoff, as indicated by the dashed line. The cutoff was determined based on the position of a substantial decrease in MeanDecreaseGini, i.e., where an “elbow” occurred in the ranking. The actual thresholds (vertical dash lines) are 5, 2, 1.5, 2, 1.2, and 1.5 for (A) to (F), respectively.

**Fig 8.**
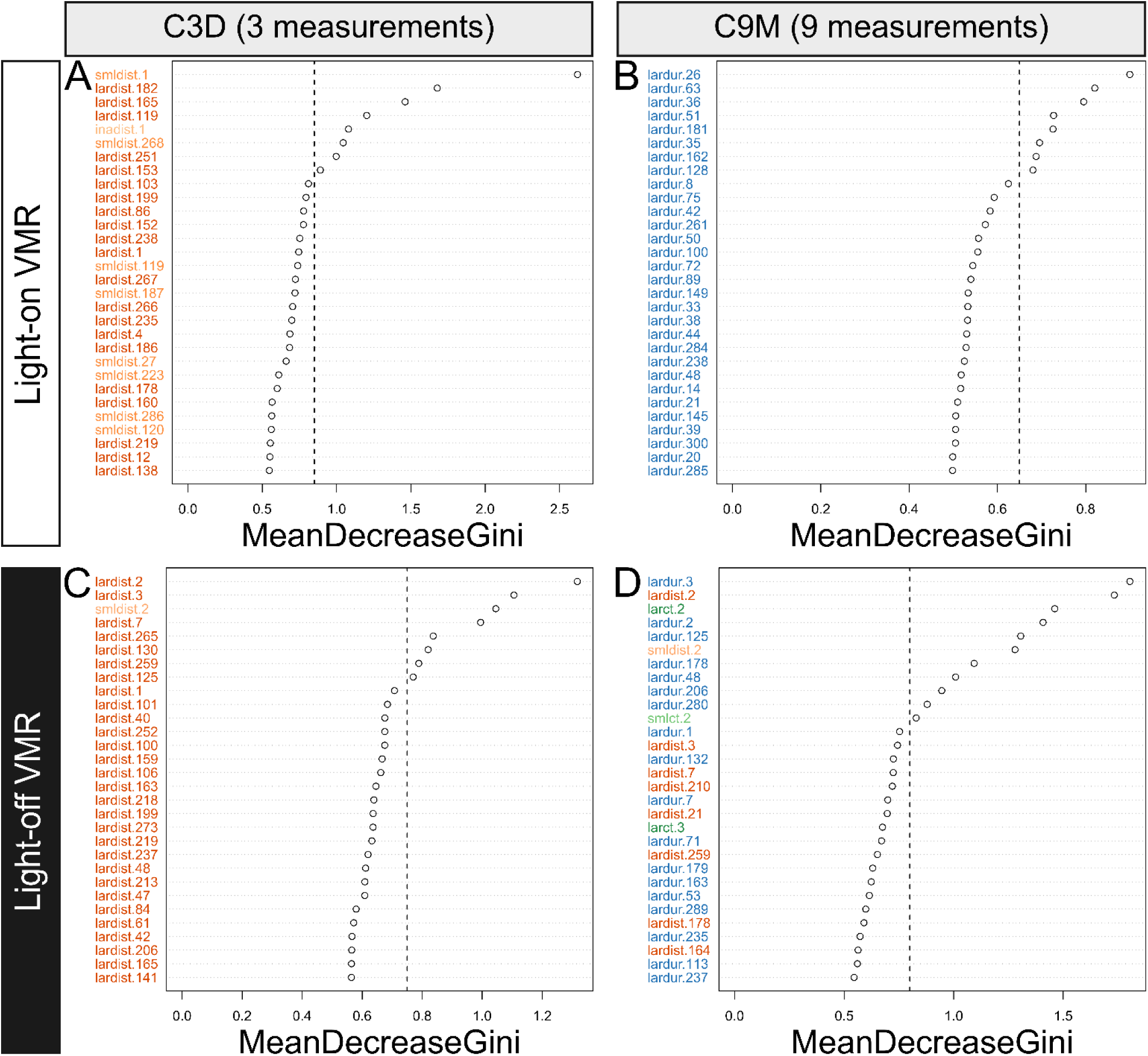
Using the embedded method to select features in the VMR datasets matricized by Feature Concatenation. (A–B) Top 30 features from the light-on dataset matricized by C3D and C9M. **(C–D)** Top 30 features from the light-off dataset matricized by C3D and C9M. These features (Y-axis) were named as in Fig 6 and ranked by the Random Forest mechanism using the mean decrease of the Gini index (MeanDecreaseGini; X-axis) in descending order as in Fig. 7. The features were color-coded based on the color scheme of features shown in Fig 2A. The selection cutoff thresholds (vertical dash lines) are 0.85, 0.65, 0.75, and 0.8 for (A) to (D), respectively.

Fig 8 shows the feature-selection results of Feature Concatenation by two approaches: C3D and C9M. In the light-on VMR data, the embedded method selected 8 out of the 900 features set from C3D (Fig 8A). Among these 8 features, *smldist.1* had the highest MeanDecreaseGini value. The embedded method selected 3 out of the 2700 features from C9D (Fig 8B), namely *lardur.164*, *lardur.48*, and *lardur.106*. In the light-off VMR data, the embedded method selected 7 out of the 900 features from C3D (Fig 8C). Among these 7 features, *lardist.2*, *lardist.3,* and *lardist.7* had the highest MeanDecreaseGini values. The embedded method also selected 6 out of 2700 features from C9M (Fig 8D). Among these 6 features, *lardur.2*, *larct.2*, and *lardur.3* had the highest MeanDecreaseGini values.

### Aggregation of selected 2DT features and performance evaluation

Table 2 lists the number of 2DT features selected by the filter and embedded methods, along with their intersection and union sets. It also includes the feature reduction rates for each selection method. In general, the number of features was reduced the most when they were selected by the intersection operation, regardless of the dataset (i.e., light-on/light-off). However, in this operation, no features were selected from the L1 Norm and L2 Norm indices and C9M for the light-on dataset. Even though this operation had the highest reduction rate (100%) for the corresponding matricization, it would not provide any feature for downstream analyses. Hence, the operation with the minimum non-zero number of features was used in the next analyses.

**Table 2.**
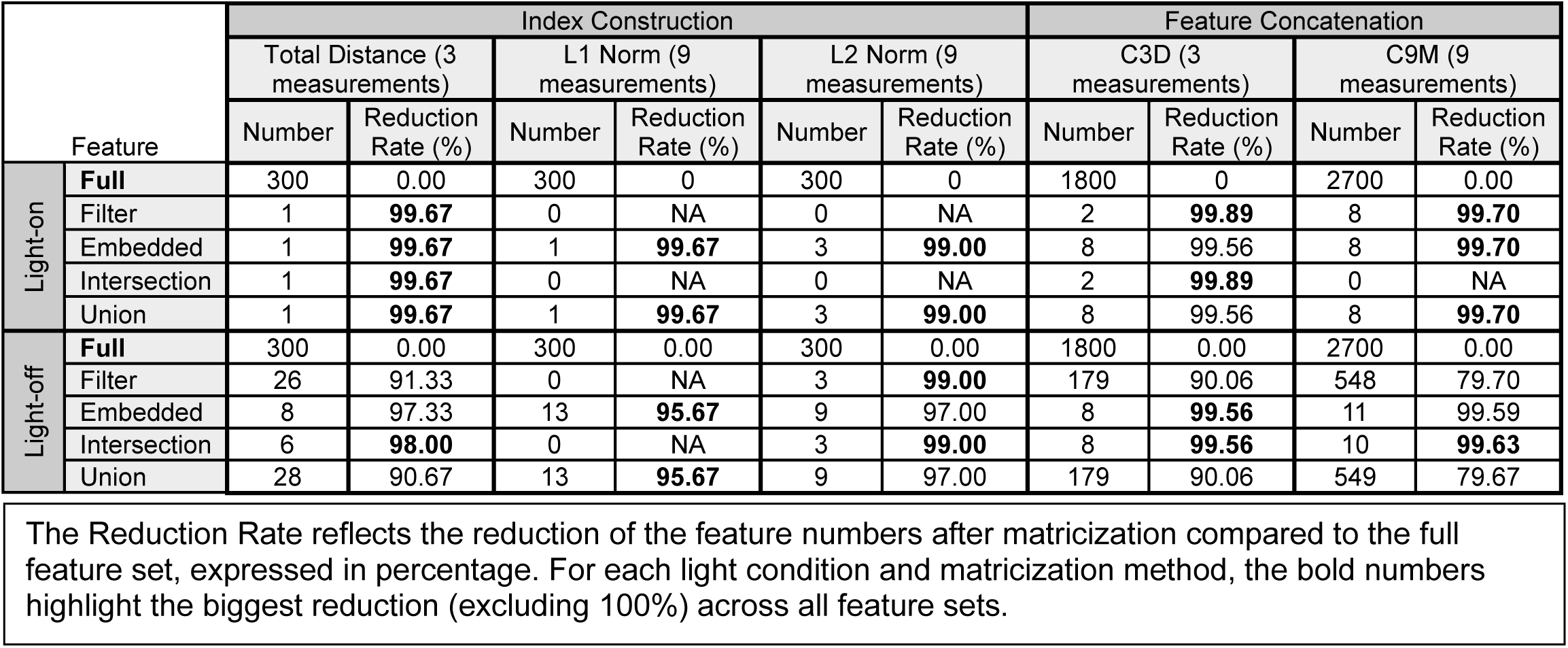
The number of features selected by different approaches.

To evaluate the impact of feature selection on MVA performance, the selected 2DT features were used in a classification task to distinguish the WT and Q344X larvae by SVM, NB, KNN, DT, RF, XGB, and MLP. A CV was conducted on the training set with 80% of the entire data, using the features selected from three constructed indices (Total Distance, L1, and L2 Norm) and two concatenated features (C3D and C9M). The resulting average AUROC for each classifier is listed in Table 3. The performance was then evaluated by comparing the highest AUROC achieved by each classifier using features from data matricized by different approaches.

**Table 3.**
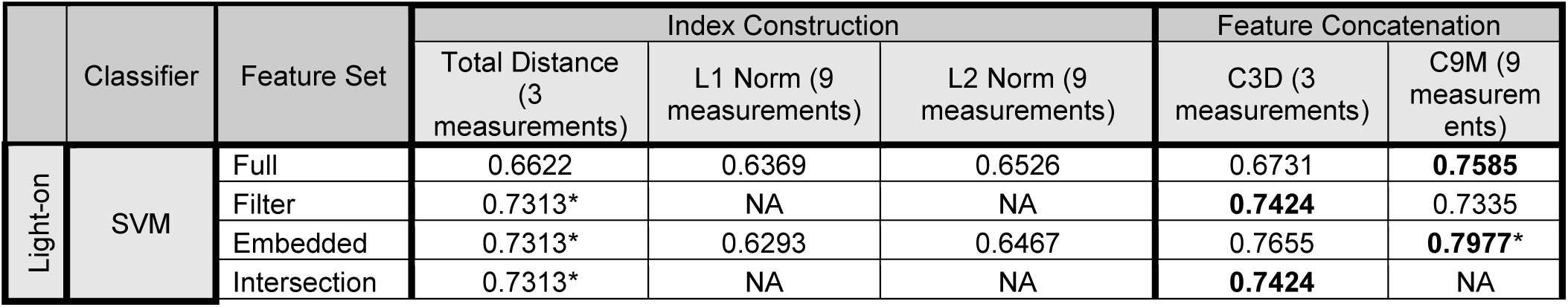
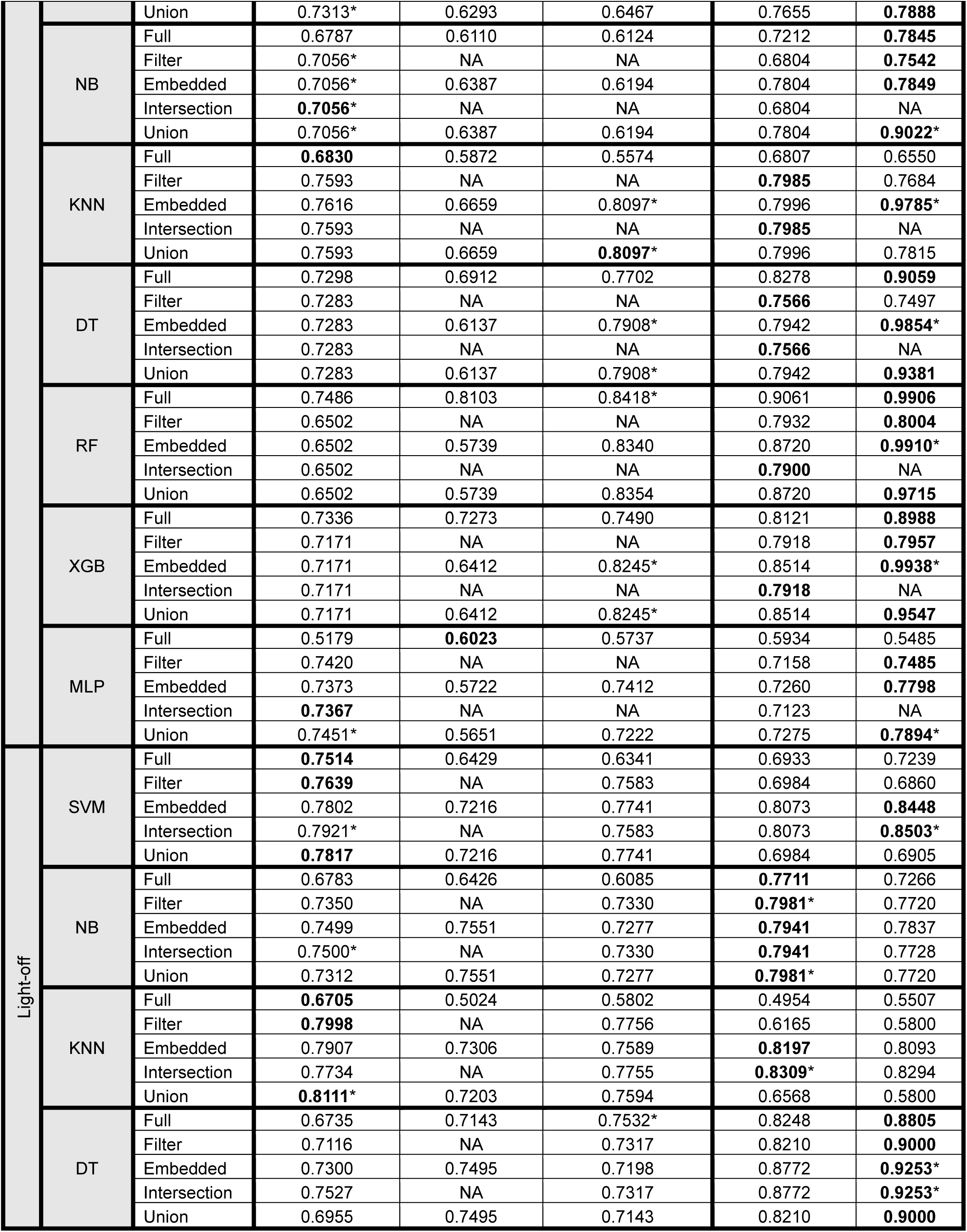
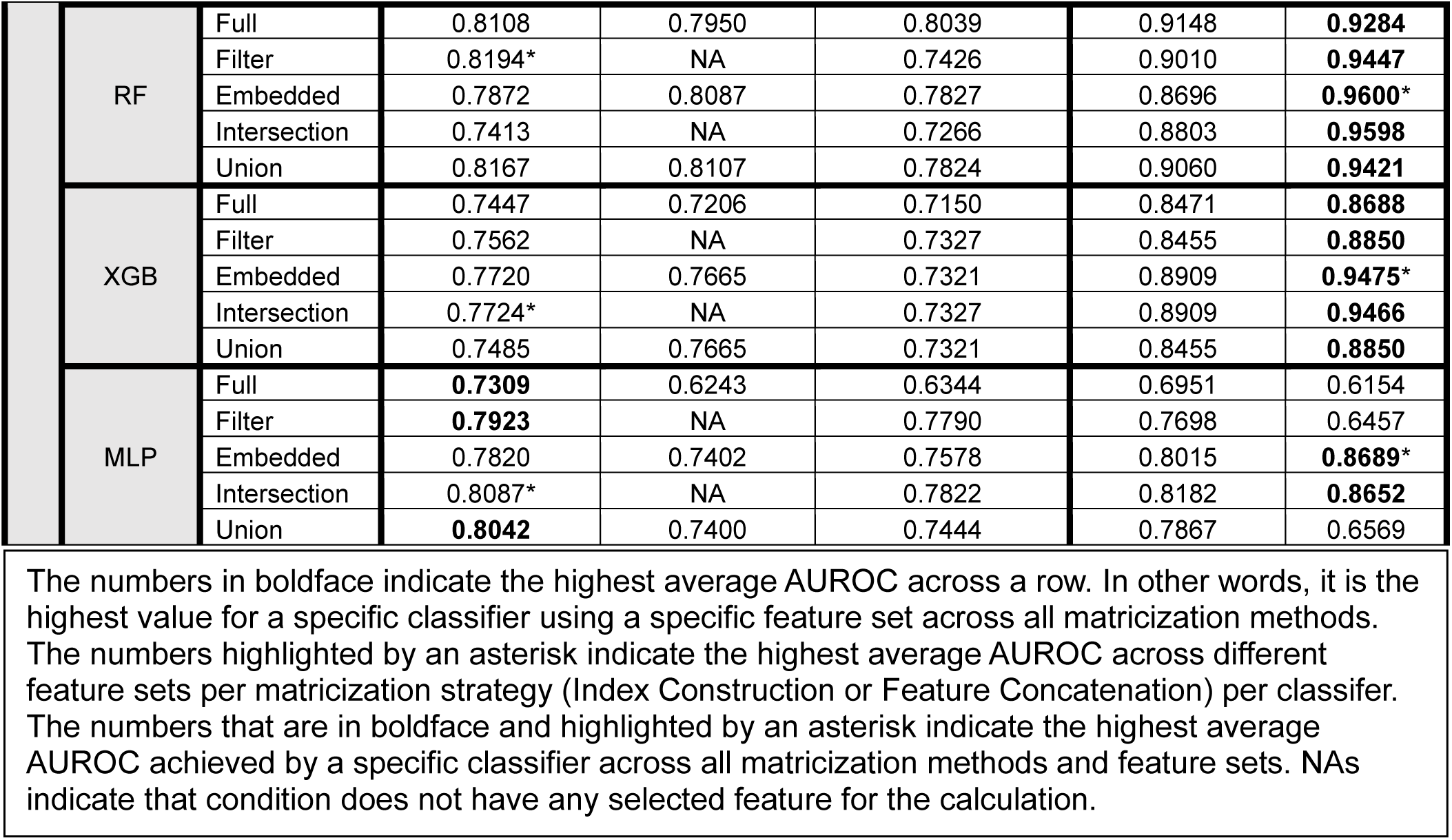
Average AUROC of CV by different classifiers using different 2DT feature sets from the data matricized by Index Construction and Feature Concatenation.

First, for the 2DT features selected from the light-on data matricized by Index Construction, 3 out of 7 classifiers achieved the highest average AUROC using the features selected from the Total Distance index (asterisks in the “Index Construction” columns, Table 3), namely SVM (0.7313), NB (0.7056), and MLP (0.7451); 3 out of 7 classifiers achieved the highest average AUROC using the features selected from the L2 Norm index, namely KNN (0.8097), DT (0.7908), and XGB(0.8245); 1 out of 7 classifiers achieved the highest average AUROC using the full feature set from the L2 Norm, namely RF (0.8418). Among the 6 classifiers that achieved the highest AUROC using selected features, 3 classifiers used either the embedded method or the union operation, namely, KNN (0.8097) and DT (0.7908); 2 classifiers used any selected feature set, namely SVM (0.7313) and NB (0.7056); MLP (0.7451) used the union operation. For the 2DT features selected from the light-off data matricized by Index Construction, 6 out of 7 classifiers achieved the highest AUROC using the features selected from the Total Distance index. Among these 6 classifiers, SVM (0.7921), NB (0.7500), XGB (0.7724), and MLP (0.8087) used the features selected by the intersection operation; KNN (0.8111) used the features selected by the union operation; RF (0.8194) used the features selected by the filter method. Overall, most classifiers achieved the highest AUROC when using features selected from the data matricized by the Total Distance index.

Second, for the 2DT features selected from the light-on data matricized by Feature Concatenation, all 7 classifiers achieved higher AUROC using the features selected from C9M than those selected from C3D (asterisks in the “Feature Concatenation” columns, Table 3).

Among them, SVM (0.7977), KNN (0.9785), DT (0.9854), RF (0.9910), and XGB (0.9938) used the features selected by the embedded method; NB (0.9022) and MLP (0.7894) used the features selected by the union operation. For the 2DT features selected from the light-off data matricized by Feature Concatenation, 5 out of 7 classifiers achieved the highest AUROC using the features selected from C9M rather than those selected from C3D. Among the 5 classifiers, RF (0.9600), XGB (0.9475), and MLP (0.8689) achieved the highest AUROC using the embedded method; DT (0.9253) achieved the highest AUROC using the embedded method and the intersection operation. Two classifiers achieved the highest AUROC using the features selected from C3D than those selected from C9M, namely NB (0.7981) and KNN (0.8309); NB used the features selected by the filter method and the union operation, whereas KNN used the features selected by the intersection operation. Overall, most classifiers achieved the highest AUROC when using features selected from the data matricized by C9M.

Finally, to evaluate the performance across both matricization methods, we compared the AUROC of different classifiers using different feature sets selected by the Total Distance index and C9M, as these AUROC tend to be the highest in their respective matricization method. In the light-on VMR data, all 7 classifiers had higher AUROC using the features from C9M (boldfaced values with asterisks, Table 3), regardless of the feature set. Among them, SVM (0.7977), KNN (0.9785), DT (0.9854), RF (0.9910), and XGB (0.9938) used the features from the embedded set; NB (0.9022) and MLP (0.7894) used the features selected by the union operation. In the light-off data, 5 out of 7 classifiers had higher AUROC using the features from C9M (boldfaced values with asterisks, Table 3). Among these 5 classifiers, SVM (0.8503), RF (0.9600), XGB (0.9475), and MLP (0.8689) used features selected by the embedded method; DT (0.9253) used features selected by the embedded method or the intersection operation. Overall, most classifiers achieved higher average AUROC when using features selected from the data matricized by Feature Concatenation rather than Index Construction.

### Generalization of the matricization approach and selected feature set to new data

After identifying the best-performing matricization approach and the resulting selected 2DT feature set in the CV, we then evaluated the extent to which their performances were applied to new data by HV (Fig 3). We first trained all classifiers using the training dataset (80% of the entire data) and all feature sets from all matricization methods, applied the optimal hyperparameters found in CV, and assessed their performances on the testing dataset (20% of the entire data). Table 4 lists the AUROC achieved by all classifiers using all feature sets selected by the Total Distance Index and C9M because they were the best-performing methods for Index Construction and Feature Concatenation, respectively. The detailed performance of all Index Construction and Feature Concatenation methods can be found in the S3 Table.

**Table 4.**
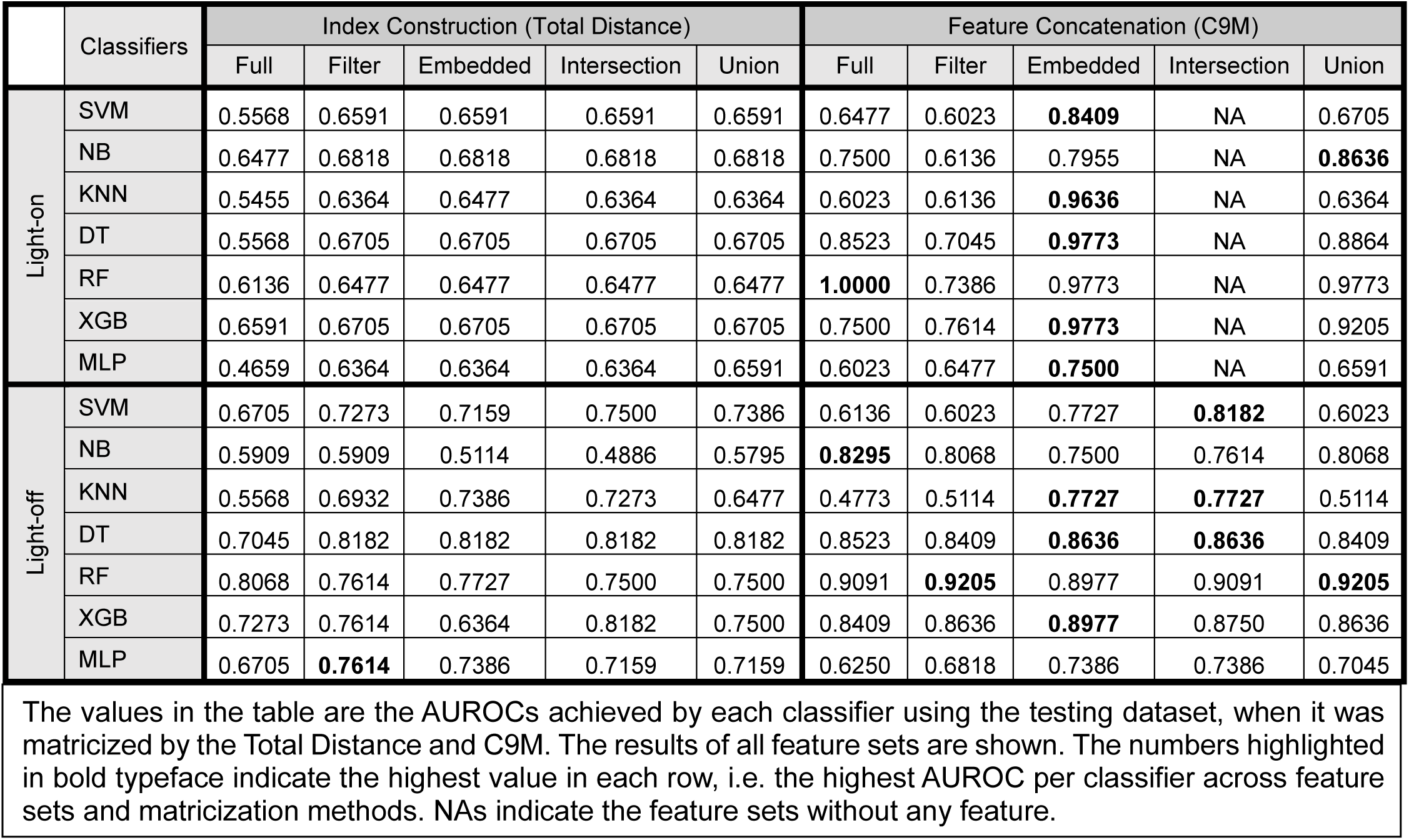
The highest AUROC of HV for Index Construction and Feature Concatenation.

|  | Classifiers | Index Construction (Total Distance) |  |  |  |  | Feature Concatenation (C9M) |  |  |  |  |
| --- | --- | --- | --- | --- | --- | --- | --- | --- | --- | --- | --- |
|  |  | Full | Filter | Embedded | Intersection | Union | Full | Filter | Embedded | Intersection | Union |
| Light-on | SVM | 0.5568 | 0.6591 | 0.6591 | 0.6591 | 0.6591 | 0.6477 | 0.6023 | **0.8409** | NA | 0.6705 |
|  | NB | 0.6477 | 0.6818 | 0.6818 | 0.6818 | 0.6818 | 0.7500 | 0.6136 | 0.7955 | NA | **0.8636** |
|  | KNN | 0.5455 | 0.6364 | 0.6477 | 0.6364 | 0.6364 | 0.6023 | 0.6136 | **0.9636** | NA | 0.6364 |
|  | DT | 0.5568 | 0.6705 | 0.6705 | 0.6705 | 0.6705 | 0.8523 | 0.7045 | **0.9773** | NA | 0.8864 |
|  | RF | 0.6136 | 0.6477 | 0.6477 | 0.6477 | 0.6477 | **1.0000** | 0.7386 | 0.9773 | NA | 0.9773 |
|  | XGB | 0.6591 | 0.6705 | 0.6705 | 0.6705 | 0.6705 | 0.7500 | 0.7614 | **0.9773** | NA | 0.9205 |
|  | MLP | 0.4659 | 0.6364 | 0.6364 | 0.6364 | 0.6591 | 0.6023 | 0.6477 | **0.7500** | NA | 0.6591 |
| Light-off | SVM | 0.6705 | 0.7273 | 0.7159 | 0.7500 | 0.7386 | 0.6136 | 0.6023 | 0.7727 | **0.8182** | 0.6023 |
|  | NB | 0.5909 | 0.5909 | 0.5114 | 0.4886 | 0.5795 | **0.8295** | 0.8068 | 0.7500 | 0.7614 | 0.8068 |
|  | KNN | 0.5568 | 0.6932 | 0.7386 | 0.7273 | 0.6477 | 0.4773 | 0.5114 | **0.7727** | **0.7727** | 0.5114 |
|  | DT | 0.7045 | 0.8182 | 0.8182 | 0.8182 | 0.8182 | 0.8523 | 0.8409 | **0.8636** | **0.8636** | 0.8409 |
|  | RF | 0.8068 | 0.7614 | 0.7727 | 0.7500 | 0.7500 | 0.9091 | **0.9205** | 0.8977 | 0.9091 | **0.9205** |
|  | XGB | 0.7273 | 0.7614 | 0.6364 | 0.8182 | 0.7500 | 0.8409 | 0.8636 | **0.8977** | 0.8750 | 0.8636 |
|  | MLP | 0.6705 | **0.7614** | 0.7386 | 0.7159 | 0.7159 | 0.6250 | 0.6818 | 0.7386 | 0.7386 | 0.7045 |

The values in the table are the AUROCs achieved by each classifier using the testing dataset, when it was matricized by the Total Distance and C9M. The results of all feature sets are shown. The numbers highlighted in bold typeface indicate the highest value in each row, i.e. the highest AUROC per classifier across feature sets and matricization methods. NAs indicate the feature sets without any feature.

In the light-on testing dataset, all 6 out of 7 classifiers achieved the highest AUROC using selected feature when the data was matricized by Feature Concatenation (boldfaced values, Table 4). Among the 6 classifiers, SVM (0.8409), KNN (0.9636), DT (0.9773), XGB (0.9773), and MLP (0.7500) used the features selected by the embedded method; NB (0.8636) used the features selected by the union operation. The only exception was RF (0.9886), which used the full feature set. In the light-off testing dataset, 5 out of 7 classifiers achieved the highest AUROC using the selected features when the data was matricized by Index Construction. Among the 5 classifiers, KNN (0.7727) and DT (0.8636) achieved the highest AUROC using the features selected by the embedded method or the intersection operation; SVM (0.8182) achieved the highest AUROC using the features selected by the intersection operation; RF (0.9205) achieved the highest AUROC using the features selected by the filter method or the union operation; XGB (0.8977) achieved the highest AUROC using the features selected by the embedded method. There were two exceptions: NB (0.8295) achieved the highest AUROC using the full feature set, and MLP (0.7614) achieved the highest AUROC using the features selected by the filter method when the data was matricized by Index Construction. Overall, most classifiers achieved the highest AUROC using the features selected by the embedded method from C9M.

### Performance evaluation of the matricization approach and selected features by other performance metrics

In addition to AUROC, other metrics can be used to evaluate performance (Table 5). These metrics were used to evaluate the extent to which the performance conclusions drawn from the AUROC metric could be generalized to other performance metrics. Table 5 lists the values of various performance metrics for different classifiers, calculated using the feature sets from C9M that had the highest AUROC for the corresponding classifier listed in Table 4.

**Table 5.** Performance evaluation by different metrics in HV using the best-performing feature set from C9M.

|  | Classifiers | AUROC | Accuracy | Sensitivity | Specificity | Precision | Kappa |
| --- | --- | --- | --- | --- | --- | --- | --- |
| Light-On | SVM | 0.8409 | 0.8409 | 0.9545 | 0.7273 | 0.7778 | 0.6818 |
|  | NB | 0.8636 | 0.8636 | 0.8636 | 0.8636 | 0.8636 | 0.7273 |
|  | KNN | 0.9636 | 0.9636 | 0.9386 | 0.9886 | 0.9880 | 0.9273 |
|  | DT | 0.9773 | 0.9773 | **1.0000** | 0.9545 | 0.9565 | 0.9545 |
|  | RF | **1.0000** | **1.0000** | **1.0000** | **1.0000** | **1.0000** | **1.0000** |
|  | XGB | 0.9773 | 0.9773 | **1.0000** | 0.9545 | 0.9565 | 0.9545 |
|  | MLP | 0.7500 | 0.7500 | 0.8636 | 0.6364 | 0.7037 | 0.5000 |
| Light-Off | SVM | 0.8182 | 0.8182 | 0.7727 | 0.8636 | 0.8500 | 0.6364 |
|  | NB | 0.8295 | 0.8295 | 0.8636 | 0.7955 | 0.8085 | 0.6591 |
|  | KNN | 0.7727 | 0.7727 | 0.6818 | 0.8636 | 0.8333 | 0.5455 |
|  | DT | 0.8636 | 0.8636 | 0.7955 | 0.9318 | 0.9211 | 0.7273 |
|  | RF | **0.9773** | **0.9773** | **1.0000** | **0.9545** | **0.9565** | **0.9545** |
|  | XGB | 0.8977 | 0.8977 | 0.8409 | **0.9545** | 0.9487 | 0.7955 |
|  | MLP | 0.7386 | 0.7386 | 0.6591 | 0.8182 | 0.7838 | 0.4773 |

In the light-on dataset, the classifier with the highest AUROC was RF (1.0000; Table 5). It also achieved the highest values in all other performance metrics. In addition, DT and XGB also achieved the same highest sensitivity (1.000 for both) as RF. In the light-off dataset, the classifiers with the highest AUROC were also RF (0.9773). It achieved the highest values in all other performance metrics. In addition, XGB also achieved the same highest specificity (0.9545) as RF. Therefore, the performance evaluated by other metrics was consistent with that evaluated by the AUROC. The conclusions drawn by AUROC can be generalized to other performance metrics.

## Discussions

Contemporary neurobehavior research often analyzes multiple behavioral measurements collected from multiple samples over time. This results in MDT data that pose challenges to the classic MVAs, as these methods are designed to handle 2DT but not MDT data. Even though MDT data can be analyzed by TCA, the resulting decomposed factors can be hard to interpret. Moreover, these results cannot be directly used for MVAs, which is the preferred analysis approach of many neurobiologists.

Consequently, they often transform the collected MDT data to 2DT data by various matricization strategies to enable the use of MVA and to ensure the interpretability of the analysis results. Various matricization methods can be implemented to transform MDT features into the simplified 2DT space. However, some 2DT features are non-informative, and some may lose useful information during the matriciztion from the original data. These 2DT features must be carefully analyzed and selected for downstream MVA to optimize the analysis performance and facilitate the interpretation of underlying behavior dynamics. In this study, we assessed different matricization and feature-selection approaches and evaluated their impacts on the performance of an MVA.

Our study utilized a 3DT behavior dataset that contained zebrafish VMR collected from the WT and visually-impaired Q344X larvae. The VMR measures several aspects of swimming activity under drastic light change, including *dur, ct,* and *dist,* at different activity levels. We first illustrated the utility of matricization in interpretation by using Index Construction to create a Total Distance index, a commonly used approach to plot and visualize VMR data [23,72,73].

The resulting plots of our VMR dataset (Fig 4) show a distinctive difference between the VMR of WT and Q344X, indicative of their difference in visual performance. In contrast, TCA only reveals some patterns in each rank of respective factors (S1 Fig), but they do not easily facilitate the interpretation of the difference between the WT and Q344X VMR. We then evaluated the impact on MVA performance by various matricization and feature-selection approaches. Specifically, we conducted a classification task to distinguish the WT and Q334X larvae using matricized data. Several classifiers were built, and CV and HV were used to evaluate the performance of different matricization and selected 2DT features. Our results indicate that most classifiers achieved the highest AUROC in the CV, using features concatenated by C9M and selected by the embedded method or the intersection operation (Table 3). Their superior performance was consistently seen in HV (Table 4) and when evaluated with other metrics (Table 5), whose performance might vary [74,75]. Their consistently high performance suggests that this specific combination of matricization and feature selection (i.e., C9M and embedded/intersection) provides the best discriminatory power in general for the VMR data. As concatenating several measurements by C9M consistently performed better than other matricization methods, this observation also indicates that different behavioral measurements in the MDT dataset provide unique information and should be included in the downstream analysis whenever appropriate. In addition, among all classifiers tested, tree-based methods (DT, RF, and XGB) achieved the highest values across various performance metrics (Table 6). This result suggested that tree-based classifiers outperformed other types of classifiers. This finding aligns with findings from benchmark studies using other types of tabular data [69,74]. Therefore, these tree-methods should be the primary choice in related VMR data analyses. Notwithstanding the general trends, there are unique differences in the classification performance using light-on and light-off datasets, particularly during the CV (Table 3). For example, RF performed the best using the full feature sets of C9M in the light-on data, while all other classifiers performed the best using either the embedded or the union feature set. In the light-off data, most classifiers, including SVM, RF, XGB, and MLP, performed the best using the embedded feature set. Overall, the embedded feature selection method and tree-based classifiers should first be considered when analyzing VMR data with similar data structures.

The selected 2DT features also reveal unique behavior dynamics between WT and Q344X larvae during light onset and offset. In the light-on VMR data, the filter method selected *inadist.1* and *smldist.1* from the C3D (Fig 6A), components of the *distance.1* feature in the Total Distance index (Fig 5A). It also selected *inadist.1* and *inadur.1* from the C9M (Fig 6B). The embedded method also consistently selected features in the first second across all indices from Index Construction (Fig 7A–C) and C3D (Fig 8A). These results indicate that the most distinctive differences in the light-on VMR of the Q344X mutant were the lack of behavior and/or a substantial reduction in the low-speed and acute response compared to the WT. At the same time, the embedded method did not select any early features from C9M; all three selected features were *lardur* in the later seconds (Fig 8B). Even in the C3D, this method selected seven *lardist* features in the later seconds out of all eight selected features (Fig 8A). These selected feature sets provided a better classification performance in CV (Table 3) and HV (Table 4). These observations indicate that although larval activities in the later seconds after light onset might not appear substantial (Fig 4A), they differed consistently in distinguishing WT and Q344X larvae. Future studies of zebrafish visual mutants by scotopic light-on VMR should consider evaluating these later components to maximize the discovery of neural mechanisms underlying the behavior differences. In the light-off VMR, the filter method selected many features with large activity from the earliest seconds onwards in both data matricized by Index Construction (Fig 5D–E) and Feature Concatenation (Fig 6C & D). In contrast, the embedded methods selected more large-activity features in the early seconds but fewer features in the later seconds, compared to the filter method (Figs 7D–7F & 8C–8D). For example, the top 3 features in C3D are *lardist.2, lardist.3,* and *lardist.7* (Fig 8C), whereas the top 3 features in C9M are *lardur.2, larct.2,* and *lardur.3* (Fig 8D). Nonetheless, these results suggest that compared to the Q344X mutant, more WT larvae tended to display an acute startle response with larger displacement for longer periods of time. This is corroborated by our classification results, in which most classifiers achieved the highest AUROC using embedded sets or intersection sets in both CV (Table 3) and HV (Table 4). Together, these results suggest that 2DT features in the early seconds after the light offset provide sufficient information to differentiate the WT and Q344X VMR. This is reasonable given that the normal light-off VMR often tends to have the largest amplitude in the early seconds (Fig 4B). Thus, the selected features from these early seconds must provide a superior discriminative power. Even though the light-off VMR also tends to have a sustained response that can provide additional information, it may not be necessary for similar discriminatory analyses.

The distinctive differences between our scotopic light-on and light-off VMR indicate that these responses are likely driven by different neural circuitries. Even though these scotopic circuitries are not well characterized, more is known about the ones driving the photopic startle response. Under a light flash (light onset), zebrafish larvae display a C-start response [76,77], consisting of a bend of about 70–80 degrees at around 183 ms, driven by the Mauthner cells [76]. Under a dark flash (light offset), zebrafish larvae display an O-bend[76,77] of approximately 150 degrees at around 408 ms [76]. This O-bend is not driven by the Mauthner cells but instead requires inputs from ventromedially-located hindbrain spinal projection neurons, including RoV3, MiV1, and MiV2 [78]. It has also been recently revealed that a distributed circuitry might control the habituation of O-bend [79,80]. The VMR is also shown by others and us to have both ocular and extra-ocular inputs [54,81,82]. The former involves contributions from both cones [83] and rods [54]. Different cone types seem to control different components of the photopic VMR [83], probably through different neural circuitries, as outlined above. However, the only rod system in the retina must control the observed scotopic VMR and innervate the corresponding neural circuitries driven by the different cone types under photopic stimulation.

Despite the new analytical and biological insights gained from our analyses, our study has several limitations. First, a few selected metrics were used for the filter methods, and one mechanism, RF, was used for the embedded method to illustrate the utility of our approach.

This selection might introduce potential bias in the feature selection. More metrics and mechanisms could be used to select features to maximize the analysis performance [47]. Second, our analyses focused on matricization along the behavior-measurement dimension. Matricization, however, can be done on any dimension. For example, the time dimension can be matricized by constructing time-sensitive features [84,85], detecting time-series motifs [32,86], or calculating the area under the curve (AUC) index for each behavioral measurement in the time series [87]. Such matricization may also enhance the downstream MVA performance and should be evaluated in future studies. Third, we acknowledge that several recent studies proposed strategies to utilize tensor decomposition results for MVA. For example, one study proposed a strategy to learn latent variables from the CP decomposition and demonstrated how these variables could be used for k-means clustering [88]. Another study proposed a strategy known as tensor decomposition-based robust feature extraction and classification, where the features were extracted in decomposed tensors, and testing data were transformed and projected to these extracted features prior to classification [89]. Even though CP decomposition results are difficult to interpret, as shown by others [89,90] and us (S1 Figure), their impact on MVA performance should be compared to that of our proposed matricization approaches. Fourth, our analysis approach may not be suitable for all behavioral studies, especially when the experimental perturbation is known to affect a specific behavioral measurement out of all possible measurements. In this case, a simple univariate analysis is likely sufficient for drawing a biological conclusion. For instance, in a study of hyperactive behavior mediated by mutations in GABA_A_ α subunits, the authors evaluated the effect of mutations on a touch-evoked response in the larvae by counting the resulting number of C-bends, a well-characterized initial phase of the C-start response [91]. Even though the larval behavior was captured by a high-speed camera and presumably MDT data could be extracted and analyzed, they were not absolutely necessary for identifying the mutations in specific GABA subunits that induced the observed hyperactive behavior.

Nonetheless, even in this case, it may be beneficial to analyze additional behavior measurements other than the C-bend with our approach to maximize the chance of detecting any effects on behavior exerted by the mutations.

In conclusion, our study demonstrated the utility of different matricization approaches and feature-selection methods in analyzing MDT behavior data. Our evaluation approach can likely apply to other types of data as long as the data structure follows the MDT format. For instance, many zebrafish studies used calcium two-photon imaging to dissect neural circuitries driving several stereotypical behaviors in response to various stimuli [77–80,92,93]. The resulting data will contain multiple dimensions, including multiple behavior measurements, time points, larvae, neurons, and their fluorescence readouts. Hence, these studies may benefit from using matricization strategies and feature-selection methods to identify relationships between the neuron firing pattern and the displayed behavior dynamics. In addition, neural-behavior studies conducted with animals other than zebrafish may also benefit from our studies, as their data also tend to consist of dimensions representing different trials, neurons, and time points of neural recordings within a single animal [17,28]. In time-series imaging studies, such as longitudinal magnetic resonance imaging [94], the data may consist of dimensions representing two dimensions of an image, samples, and time points [95,96]. To analyze these MDT data, we recommend the following workflow: Step 1) Preprocess and normalize the data using the standard procedures in the field; Step 2) Transform the MDT data to lower-order tensors by matricizing the dimensions of interest with the preferred approaches; Step 3) Select features from the matricized data with different feature-selection methods; Step 4) Evaluate the performance of specific combinations of matricization approaches and feature- selection methods; and Step 5) Applied the best-performing selected features for downstream MVA. By incorporating more information into the analysis, we anticipate our workflow will uncover novel insights into complex neural behavior.

## Acknowledgments

We thank Mengrui Zhang for his thoughtful discussions.This publication was made possible, in part, with support from the Indiana Clinical and Translational Sciences Institute to YFL and BW, funded in part by Grant Number UM1TR004402 from the National Institutes of Health, National Center for Advancing Translational Sciences, Clinical and Translational Sciences Award. The content is solely the responsibility of the authors and does not necessarily represent the official views of the National Institutes of Health. JC, LF and PM were partially supported by National Science Foundation grants DMS-2124493, DMS- 2311297, DMS-2319279, DMS-2318809, and National Institutes of Health grants R01GM152814.

## Notes

https://doi.org/10.7910/DVN/B8HBU9

### Competing Interest Statement

The authors have declared no competing interest.

