## Supplementary material for "Tensor analysis of animal behavior by matricization and feature selection": S1 Appendix

**S1 Appendix. Data normalization**

Prior to matricization, the raw VMR data were normalized against several confounding factors, including light-intensity variations across the locations in a 96-well plate, batch effect, and baseline activity variations across independent experiments [1]. The normalization involves fitting the following linear regression model:

$$y{}_{ij}=\boldsymbol{x}_{ij}^{T}\boldsymbol{\beta}_{j}\boldsymbol{+}\epsilon_{ij}$$

where *y_ij_* denotes the value of a behavioral measurement at a second for $i$th larvae in group *j* (i.e., different locations in the 96-well plate, different batches, different genotypes/treatments); ***x****_ij_* denotes a column vector of explanatory variables for the corresponding larva; ***β****_j_* denotes a column vector of coefficients for the explanatory variables in the linear regression model; *ϵ_ij_* represents the residuals. Note in the baseline activity normalization, only ***β****_j_* was present in the equation and estimated by the average dark activity value of group *j*. In the normalization, each confounding factor was fitted to this linear regression model to calculate the residues ***ϵ****_ij_*, which was then used as the normalized data.
