## Supplementary material for "Tensor analysis of animal behavior by matricization and feature selection": S1 Figure

**S1 Figure. Tensor Component Analysis (TCA) results of the 3DT VMR data.** In TCA, each dimension was decomposed to the multiple components or ranks of the subspace to represent the data points in the original 3DT space. Here, only the first three ranks are shown. **(A–C)** The light-on VMR data decomposed into time, measurement, and sample factors. **(A)** The first three ranks of the time factor after decomposition in the light-on VMR data. Data points are tensor decomposition scores (y-axis) for each time point (x-axis). In all three components, larval movement increased over time, regardless of the genotype. **(B)** The first three ranks of the behavior measurement factor after decomposition in the light-on VMR data. The data points are tensor decomposition scores (y-axis) for each behavior measurement (x-axis). In all three components, *inact* values are much higher than other behavior measurements, regardless of the genotype. **(C)** The first three ranks of the sample factor after decomposition in the light-on VMR data. The data points are tensor decomposition scores (y-axis) for each sample (x-axis). The samples were ordered based on the genotypes, Q344X (red) and WT (black). The result shows three groups of larvae displaying distinct behaviors, but there is no difference between the two genotypes. **(D–F)** The light-off VMR data decomposed into three factors: time, measurement, and sample factors. For each factor, only the first three ranks were shown. **(D)** The first three ranks of the time factor after decomposition in the light-off VMR data. The data points are tensor decomposition scores (y-axis) for each time point (x-axis). There are three trends of behavior values, regardless of the genotype: Rank 1 shows a trend where larvae slowly become more active; Rank 2 shows a trend where larvae suddenly become active and stay active; Rank 3 shows a trend where larvae are not super active in the beginning but slowly becomes more active, and later quiets down a bit. **(E)** The first three ranks of the behavior measurement factor after decomposition in the light-off VMR data. Data points are tensor decomposition scores (y-axis) for each behavior measurement (x-axis). Different measurements have different scores. The trends are the same in all three ranks, that *inact* and *smlct* have much higher values than the rest of the measurements, regardless of the genotype. **(F)** The first three ranks of the sample factor after tensor decomposition in the light-off VMR data. The data points are tensor decomposition scores (y-axis) for each sample (x-axis). The samples were ordered based on the genotype Q344X (red) and WT (black). There are subtle differences between WT and Q344X: In Rank 1, WT and Q344X had similar distributions on the score; In Rank 2, Q344X had lower average scores than WT; In Ranks 3, Q344X were more dispersed along the Score values than WT.

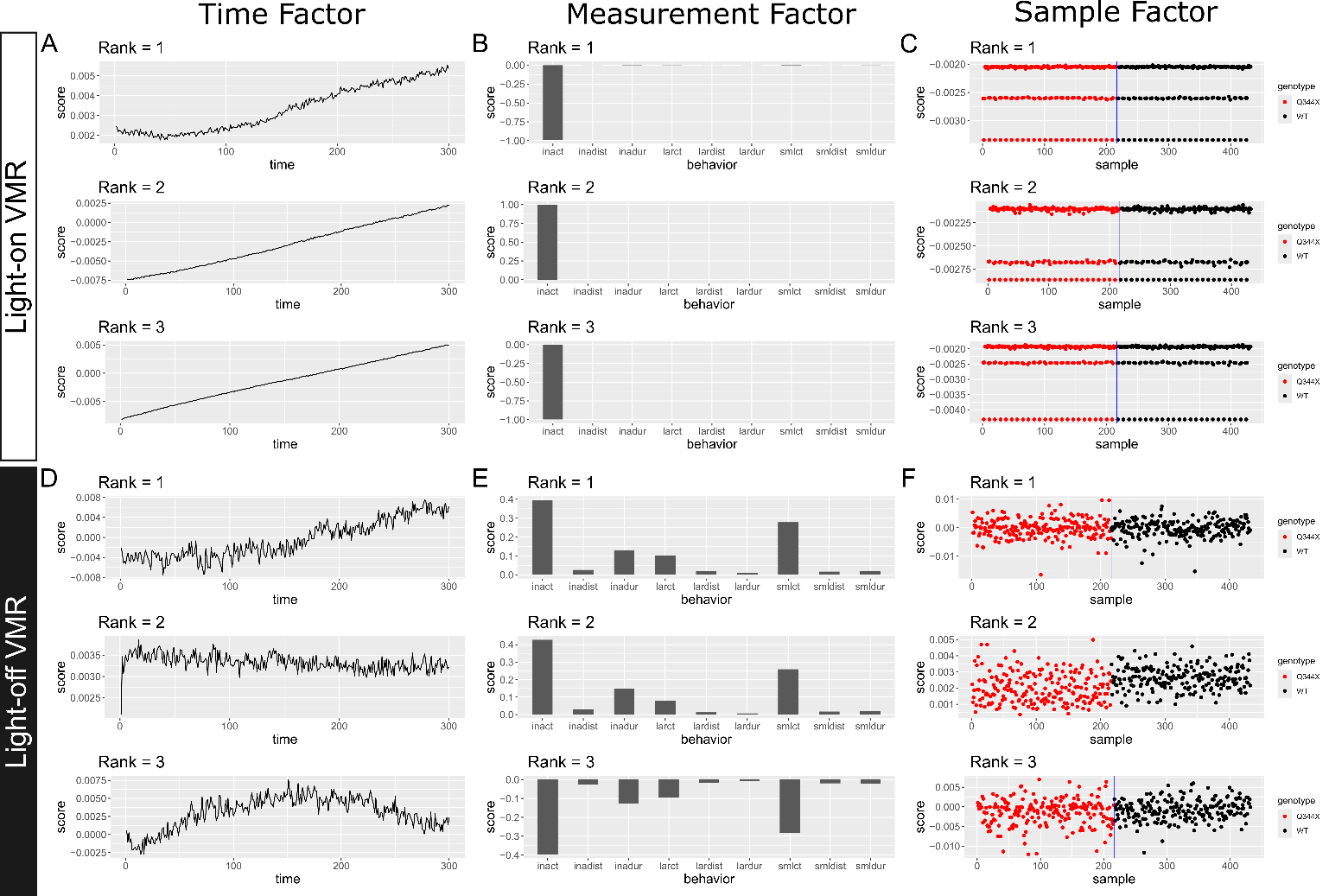
