## Supplementary material for "Tensor analysis of animal behavior by matricization and feature selection": S1 Table

**S1 Table.** **Names and descriptions of behavior measurements recorded in the zebrafish VMR data.**

| Short name | Long name | Description |
| --- | --- | --- |
| *inadur* | inactivity duration | The time the larva spends swimming at the inactivity speed (< 0.6 cm/s) within each time bin. |
| *smldur* | small activity duration | The time the larva spends swimming at the small activity (≥ 0.6 cm/s and < 1 cm/s) within each time bin. |
| *lardur* | large activity duration | The time the larva spends swimming at the large activity speed (≥ 1 cm/s) within a time bin. |
| *inact* | inactivity count | The number of times the larva movement speed crosses the threshold and enters the inactivity speed range (< 0.6 cm/s) within a time bin. |
| *smlct* | small activity count | The number of times the larva movement speed crosses the threshold and enters the small speed range (≥ 0.6 cm/s and < 1 cm/s) within a time bin. |
| *larct* | large activity count | The number of times the larva movement speed crosses the threshold and enters the large activity speed range (≥ 1 cm/s) within a time bin. |
| *inadist* | inactivity distance | The distance the larva travels when swimming at the inactivity speed (< 0.6 cm/s) within each time bin. |
| *smldist* | small activity distance | The distance the larva travels when swimming at the small activity speed (≥ 0.6 cm/s and < 1 cm/s) within each time bin. |
| *lardist* | large activity distance | The distance the larva travels when swimming at the large activity speed (≥ 1 cm/s) within each time bin. |
