## Supplementary material for "Tensor analysis of animal behavior by matricization and feature selection": S2 Table

**S2 Table.** **Package information and hyperparameter tuning space for classifiers used in this study.** * Applicable when kernel = polynomial, radial, or sigmoid. † Only when kernel = polynomial. ‡ Only when kernel = polynomial or sigmoid.

| Classifiers | Language | Package | Hyperparameter Range |
| --- | --- | --- | --- |
| SVM | R | *e1071*, Version = 1.7-13 | **kernel**: Linear, Polynomial, Radial, Sigmoid  **gamma***: 1, 2,3  **degree**†: 2,3  **coefficient**‡: 0, 1, 2, 3, 4, 5 |
| NB | R | *naivebayes*, Version = 0.9.7 | **Usekernel**: True, False  **laplace**: 0,1,2,3 |
| KNN | R | *caret*, Version = 6.0-94 | **k**: 1 to 35 |
| DT | R | *rpart*, Version = 4.1-15 | **split**: Gini, Information |
| RF | R | *randomForest*, Version = 4.7-1.1 | **ntree**: 125, 250, 500, 1000  **mtry**: 8, 16, 32, 64, 128, 256 |
| XGB | R | *xgboost*, Version = 1.7.3.1 | **eta**: 0.5, 1, 1.5, 2  **gamma**: 0, 0.1, 0.4, 0.8  **max_depth**: 3 ,6, 9 |
| MLP | Python | Keras, version = 2.3.1 | **units**(Dense): 8, 16, 32, 64, 128  **Optimizer**: Adam  **learning_rate**(Adam): 0.0001, 0.001, 0.01  **epoch**: 500 |
