## Supplementary material for "Tensor analysis of animal behavior by matricization and feature selection": S3 Table

**S3 Table. The AUROC of holdout validation for Index Construction and Feature Concatenation.** The values in the table are the AUROC achieved by each classifier using the testing dataset when it was matricized by different Index Construction and Feature Concatenation methods. The numbers highlighted in bold typeface indicate the highest value in each row, i.e., the highest AUROC per classifier per feature set across different matricization methods. NAs indicate the feature sets without any feature. The asterisks indicate the highest AUROC per classifier per matricization method across different feature sets.

|  | Classifiers | Feature | Index Construction | | | Feature Concatenation | |
| --- | --- | --- | --- | --- | --- | --- | --- |
|  |  |  | Total Distance (3 measurements) | L1 Norm (9 measurements) | L2 Norm (9 measurements) | C3D (3 measurements) | C9M (9 measurements) |
| Light-on | SVM | full | 0.5568 | 0.5341 | 0.5909 | 0.5341 | **0.6477** |
|  |  | filter | **0.6591*** | NA | NA | 0.6364 | 0.6023 |
|  |  | embedded | 0.6591***** | 0.5568 | 0.6250 | 0.6591 | **0.8409*** |
|  |  | intersection | **0.6591*** | NA | NA | 0.6364 | NA |
|  |  | union | 0.6591***** | 0.5568 | 0.6250 | 0.6591 | **0.6705** |
|  | NB | full | 0.6477 | 0.6023 | 0.5114 | 0.6705 | **0.7500** |
|  |  | filter | **0.6818*** | NA | NA | 0.6250 | 0.6136 |
|  |  | embedded | 0.6818***** | 0.6250 | 0.5568 | 0.6591 | **0.7955** |
|  |  | intersection | **0.6818*** | NA | NA | 0.6250 | NA |
|  |  | union | 0.6818***** | 0.6250 | 0.5568 | 0.6591 | **0.8636*** |
|  | KNN | full | 0.5455 | 0.5568 | 0.5000 | 0.5341 | **0.6023** |
|  |  | filter | 0.6364 | NA | NA | **0.7045** | 0.6136 |
|  |  | embedded | 0.6477 | 0.5909 | 0.7386***** | 0.7614 | **0.9636*** |
|  |  | intersection | 0.6364 | NA | NA | **0.7045** | NA |
|  |  | union | 0.6364 | 0.5909 | 0.7386***** | **0.7614** | 0.6364 |
|  | DT | full | 0.5568 | 0.6477 | 0.6591 | 0.7841 | **0.8523** |
|  |  | filter | 0.6705 | NA | NA | 0.6705 | **0.7045** |
|  |  | embedded | 0.6705 | 0.5227 | 0.7955***** | 0.7500 | **0.9773*** |
|  |  | intersection | **0.6705** | NA | NA | **0.6705** | NA |
|  |  | union | 0.6705 | 0.5227 | 0.7955***** | 0.7500 | **0.8864** |
|  | RF | full | 0.6136 | 0.7727 | 0.7386 | 0.9091 | **1.0000*** |
|  |  | filter | 0.6477 | NA | NA | 0.7273 | **0.7386** |
|  |  | embedded | 0.6477 | 0.5455 | 0.7955 | 0.8523 | **0.9773** |
|  |  | intersection | 0.6477 | NA | NA | **0.7159** | NA |
|  |  | union | 0.6477 | 0.5455 | 0.8068***** | 0.8523 | **0.9773** |
|  | XGB | full | 0.6591 | 0.6136 | 0.7045 | 0.7045 | **0.7500** |
|  |  | filter | 0.6705 | NA | NA | 0.7045 | **0.7614** |
|  |  | embedded | 0.6705 | 0.5568 | 0.7955***** | 0.8295 | **0.9773*** |
|  |  | intersection | 0.6705 | NA | NA | **0.7045** | NA |
|  |  | union | 0.6705 | 0.5568 | 0.7955***** | 0.8295 | **0.9205** |
|  | MLP | full | 0.4659 | **0.6023** | 0.5227 | 0.5114 | **0.6023** |
|  |  | filter | 0.6364 | NA | NA | **0.6477** | **0.6477** |
|  |  | embedded | 0.6364 | 0.5682 | **0.7727*** | 0.5795 | 0.7500* |
|  |  | intersection | **0.6364** | NA | NA | 0.5682 | NA |
|  |  | union | 0.6591 | 0.6136 | **0.7045** | 0.6136 | 0.6591 |
| Light-off | SVM | full | **0.6705** | 0.5682 | 0.6591 | 0.6250 | 0.6136 |
|  |  | filter | **0.7273** | NA | **0.7273** | 0.6591 | 0.6023 |
|  |  | embedded | 0.7159 | 0.7159 | 0.6818 | **0.8182*** | 0.7727 |
|  |  | intersection | 0.7500 | NA | 0.7273 | **0.8182*** | **0.8182*** |
|  |  | union | **0.7386*** | 0.7159 | 0.6818 | 0.6591 | 0.6023 |
|  | NB | full | 0.5909 | 0.6818 | 0.5909 | 0.6932 | **0.8295*** |
|  |  | filter | 0.5909 | NA | 0.6023 | 0.6591 | **0.8068** |
|  |  | embedded | 0.5114 | 0.7045***** | 0.5795 | 0.5795 | **0.7500** |
|  |  | intersection | 0.4886 | NA | 0.6023 | 0.5795 | **0.7614** |
|  |  | union | 0.5795 | 0.7045***** | 0.5795 | 0.6591 | **0.8068** |
|  | KNN | full | **0.5568** | 0.4886 | 0.5114 | 0.4659 | 0.4773 |
|  |  | filter | 0.6932 | NA | **0.7500*** | 0.5909 | 0.5114 |
|  |  | embedded | 0.7386 | 0.6023 | 0.5455 | **0.7955** | 0.7727 |
|  |  | intersection | 0.7273 | NA | 0.7273 | **0.8068*** | 0.7727 |
|  |  | union | **0.6477** | 0.5909 | 0.5682 | 0.5568 | 0.5114 |
|  | DT | full | 0.7045 | 0.7841 | 0.6477 | 0.7841 | **0.8523** |
|  |  | filter | 0.8182***** | NA | 0.7159 | 0.7727 | **0.8409** |
|  |  | embedded | 0.8182***** | 0.7500 | 0.5568 | **0.8977*** | 0.8636 |
|  |  | intersection | 0.8182***** | NA | 0.7159 | **0.8977*** | 0.8636 |
|  |  | union | 0.8182***** | 0.7500 | 0.5568 | 0.7727 | **0.8409** |
|  | RF | full | 0.8068***** | 0.7614 | 0.7386 | **0.9091** | **0.9091** |
|  |  | filter | 0.7614 | NA | 0.7500 | 0.9091 | **0.9205*** |
|  |  | embedded | 0.7727 | 0.7614 | 0.6932 | **0.9091** | 0.8977 |
|  |  | intersection | 0.7500 | NA | 0.7614 | 0.8977 | **0.9091** |
|  |  | union | 0.7500 | 0.7614 | 0.7045 | 0.8977 | **0.9205*** |
|  | XGB | full | 0.7273 | 0.6477 | 0.7159 | **0.8409** | **0.8409** |
|  |  | filter | 0.7614 | NA | 0.7045 | 0.8295 | **0.8636** |
|  |  | embedded | 0.6364 | 0.6818 | 0.6932 | 0.8864 | **0.8977*** |
|  |  | intersection | 0.8182***** | NA | 0.7045 | **0.8864** | 0.8750 |
|  |  | union | 0.7500 | 0.6818 | 0.6932 | 0.8295 | **0.8636** |
|  | MLP | full | **0.6705** | 0.5909 | 0.5568 | 0.6477 | 0.6250 |
|  |  | filter | **0.7614*** | NA | 0.7386 | 0.6932 | 0.6818 |
|  |  | embedded | 0.7386 | 0.6364 | 0.5795 | **0.7727*** | 0.7386 |
|  |  | intersection | 0.7159 | NA | 0.6932 | **0.7500** | 0.7386 |
|  |  | union | **0.7159** | 0.6477 | 0.6591 | 0.6705 | 0.7045 |
